# Targeting of subcellular metabotropic glutamate receptor 5 signaling to modulate pain transmission

**DOI:** 10.64898/2026.01.29.702703

**Authors:** Jeffri S. Retamal, Shane D. Hellyer, Paulina D. Ramirez-Garcia, Rocco Latorre, Rina Pokhrel, Thomas P. Davis, Yifei Zhu, Michael R. Whittaker, Jackson A. Kos, Kelly O’Sullivan, Nigel W. Bunnett, Wendy L. Imlach, Daniel P. Poole, Karen J. Gregory, Nicholas A. Veldhuis

## Abstract

Metabotropic glutamate receptor 5 (mGlu_5_) is a class C GPCR crucial for neuronal development and synaptic transmission. mGlu_5_ is a potential therapeutic target in pain management and modulates pain-associated gene expression and signaling pathways. Although mGlu_5_ inhibitors have shown promise in treating pain, none have translated to the clinic. Up to 90% of neuronal mGlu_5_ expression is intracellular, although the precise locations and function of different mGlu_5_ intracellular pools remains unclear. Building on recent evidence showing the importance of endosome-mediated nociceptive signaling by other GPCRs, we hypothesized that endosomal pools of mGlu_5_ contribute to pain transmission, and that targeted inhibition of intracellular mGlu_5_ signaling results in superior analgesia. Using calcium mobilization assays and genetically encoded resonance energy transfer biosensors, we report that upon its activation mGlu_5_ recruits Gα_q/11_ and Gα_s_ to the plasma membrane. Conversely, internalized mGlu_5_ in endosomes recruits only Gα_q/11_ proteins. mGlu_5_ signaling is highly dependent on receptor trafficking to endosomes, with sustained nuclear ERK1/2 signaling requiring both receptor internalization and active glutamate transport into the cell. We generated pH responsive nanoparticles loaded with the mGlu_5_ negative allosteric modulator VU0366058 (DIPMA-VU058), enabling endosome-targeted inhibition of mGlu_5_. Nanoparticle encapsulation of VU0366058 enhanced inhibition of both acute and sustained nuclear ERK1/2 signaling, and significantly reduced neuronal excitability in nociceptive circuits in spinal cord slices from rats with neuropathic pain. Intrathecal administration of DIPMA-VU058 achieved superior analgesia in both inflammatory and neuropathic models of pain in mice compared to free VU0366058 and the reference compound fenobam. These studies demonstrate the importance of endosome-associated receptors for the complete mGlu_5_ signaling response. Furthermore, we show that manipulating the cellular distribution of an allosteric modulator can engender location-biased pharmacological effects. Together, we have revealed new and unappreciated roles for endosome-specific mGlu5 signaling and demonstrate that endosome-selective targeting may offer an alternative therapeutic approach for modulating mGlu_5_ activity.

## Introduction

Glutamate, the major excitatory neurotransmitter in the mammalian CNS, exerts its actions through ionotropic and metabotropic glutamate (mGlu) receptors^1,2^. Ionotropic receptors mediate fast synaptic transmission, with G protein-coupled mGlu receptors contributing to slower, neuromodulatory responses^1,2^. mGlu receptors are classified into three subgroups based on their G protein coupling, signaling and sequence homology; Group I (mGlu_1_ and mGlu_5_), Group II (mGlu_2_ and mGlu_3_) and Group III (mGlu_4_, mGlu_6_, mGlu_7_ and mGlu_8_)^1^. Group I receptors canonically couple to Gα_q/11_, resulting in activation of phospholipase C (PLC) and inositol phosphate mediated intracellular calcium (iCa^2+^) mobilization^3^. This is followed by activation of downstream kinases such as protein kinase C, extracellular signal regulated kinase 1/2 (ERK1/2) and pleiotropic signaling through other G proteins, downstream effectors and modulation of ion channel function^1^. mGlu_5_ is widely expressed in many brain regions and in the spinal cord ^4–9^, and is a crucial modulator of synaptic plasticity and neuronal development ^10–13^. Dysregulation of mGlu_5_ signaling is associated with multiple neurological disorders, including inflammatory, neuropathic and chronic pain conditions ^7,14–19^. As such, mGlu_5_ has gained interest as a potential target for development of novel analgesic drugs.

Up to 90% of neuronal mGlu_5_ is intracellular, particularly on nuclear and endoplasmic reticular membranes, with intracellular receptors mediating distinct signaling cascades compared to cell surface receptors ^20–25^. In spinal neurons, mGlu_5_ localizes to the endoplasmic reticulum and inner nuclear membrane, where expression is upregulated in rodent models of pain^7,14^. Enhanced nuclear mGlu_5_ activation upregulates transcription factors mediating pain behavior, with excitatory amino acid transporter 3 (EAAT3) playing an important role in supplying glutamate to mGlu_5_ at the nuclear membrane ^7,14,20,22^. In addition to intracellular mGlu_5_, cell surface mGlu_5_ internalizes both constitutively and upon activation via clathrin- and caveolar mediated endocytosis ^26–29^. Endocytosed mGlu_5_ co-localizes with markers of early and recycling endosomes, but not lysosomes, with dynamic mGlu_5_ trafficking between cell surface and intracellular compartments ^22,27,28,30–32^. Recent evidence indicates GPCRs can continue to signal from endosomes with distinct signaling and functional outcomes, including signaling related to nociception and pain ^33–44^. Importantly, selective modulation of endosomal GPCRs results in prolonged analgesia in multiple pain models ^33–37,41,45,46^. While endosomal signaling has been demonstrated for other pain-related receptors, whether mGlu_5_ induces sustained signaling from endosomes is unclear. As such, the therapeutic potential of modulating such signaling remains unknown.

Targeting intracellular mGlu_5_ with negative allosteric modulators (NAMs), small molecules that bind a site distinct from that of glutamate, reduces neuropathic pain behaviors in animal models to a greater extent than blocking cell surface signaling ^7,14^. NAMs from distinct chemical scaffolds display different cell-type and location-dependent inhibition of intracellular mGlu_5_, suggesting rational drug design or drug delivery methods to increase intracellular exposure may enhance the therapeutic potential of mGlu_5_ NAMs in treating pain ^47^. While mGlu_5_ NAMs have demonstrated promising results in animal models of pain ^7,9,14,15,48–51^, a lack of analgesic efficacy and tolerance and other adverse effects in humans has hampered translation ^52,53^. This poor clinical profile could be related to ineffective targeting of mGlu_5_ at the correct location to modulate pain signaling in humans. Here, we investigated the potential contribution of endosomal signaling to pain-relevant mGlu_5_ responses *in vitro* and *in vivo*. We showed that glutamate induces mGlu_5_ internalization into early endosomes in HEK293A cells and recruits Gα_q/11_ proteins to endosomes, accounting for a small but definable component of sustained mGlu_5_ signaling. We developed pH responsive nanoparticles to selectively target endosomes and demonstrated co-distribution with fluorescently tagged mGlu_5_. Further, we demonstrated that nanoparticles containing the mGlu_5_ NAM VU0366058 more efficiently inhibited mGlu_5_ signaling in HEK293A cells and blocked agonist-induced electrophysiological changes in spinal cord neurons. Intracellular delivery using nanoparticles resulted in more effective analgesia compared to free VU0366058 in mouse models of both inflammatory and neuropathic pain. Together, these data show that endosomal signalling is a minor but important component of overall mGlu_5_ function. Further, with nanoparticle-mediated targeting of an allosteric modulator to intracellular mGlu_5_ pools, we demonstrate that influencing the location-biased properties of mGlu_5_ NAMs is a new and viable method for enhancing their analgesic efficacy.

## KEY RESOURCES TABLE

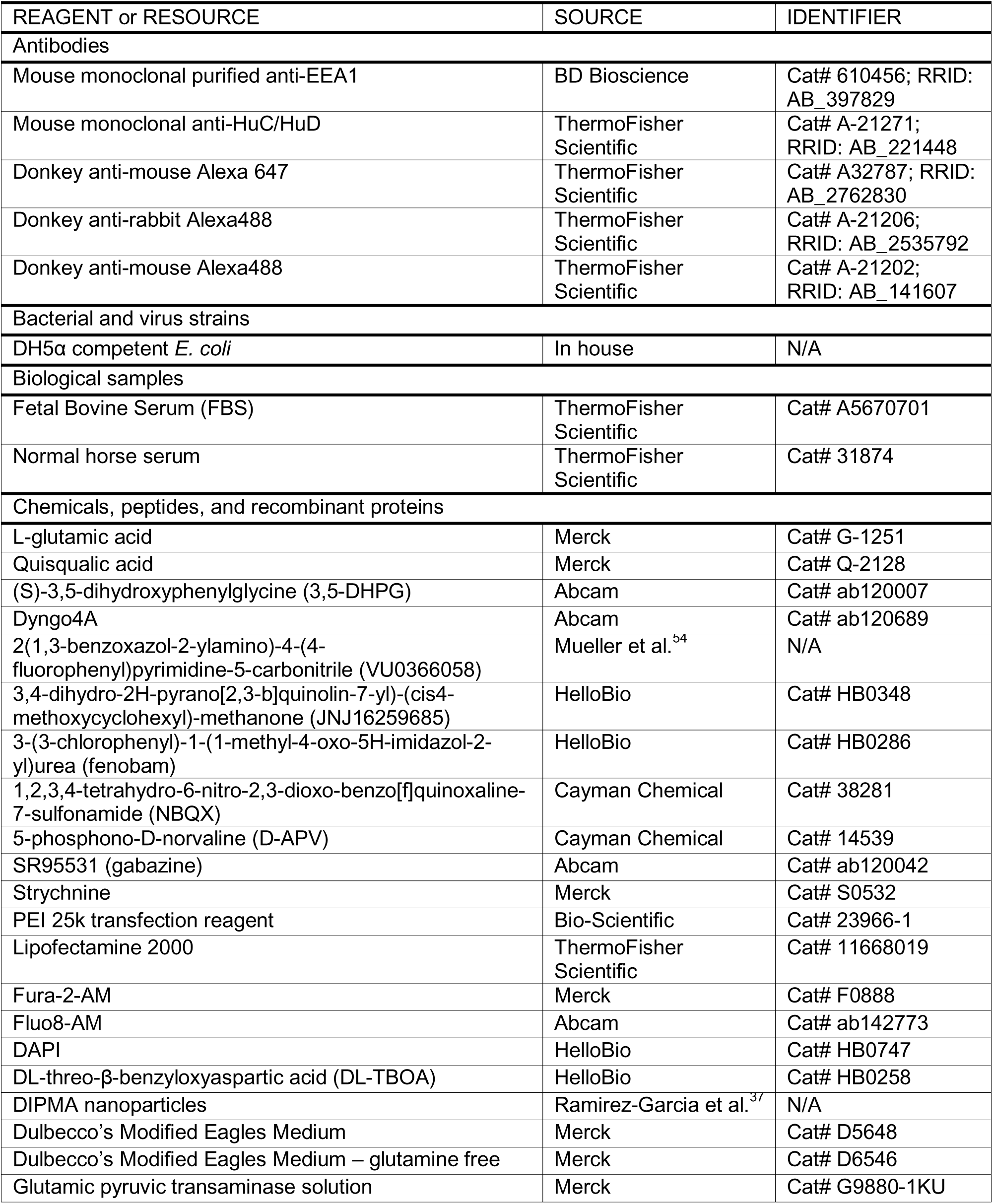
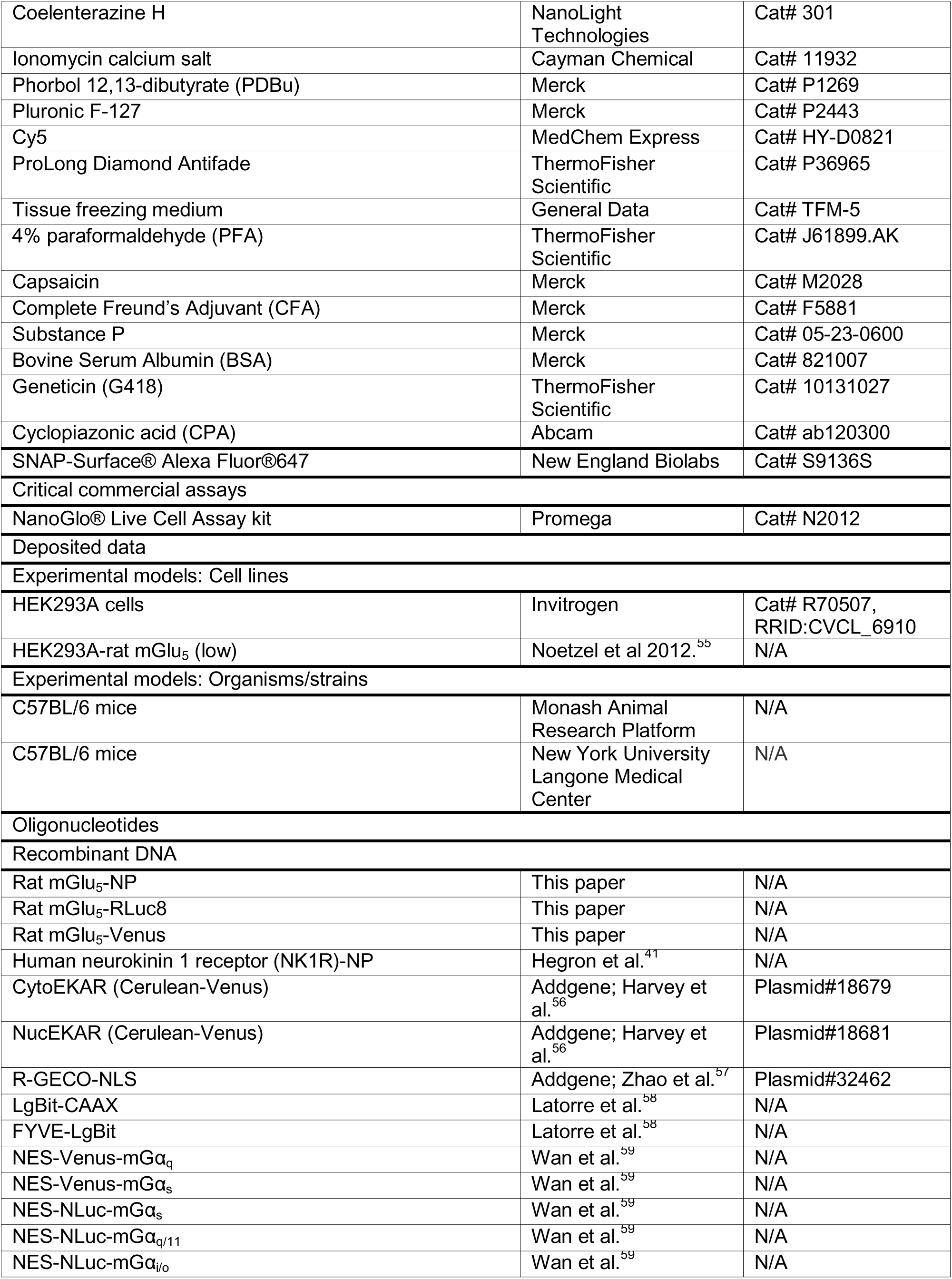
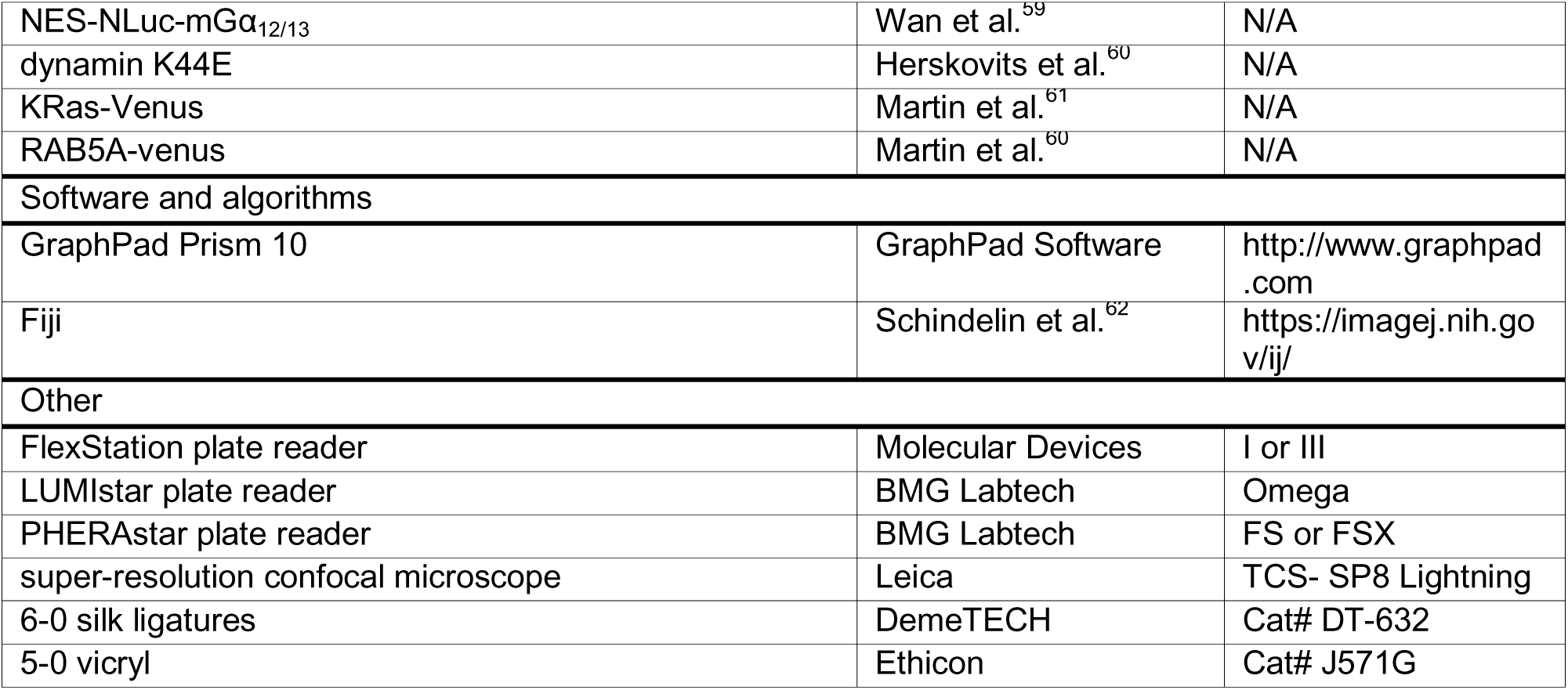

## RESOURCE AVAILABILITY

### Lead contact

Further information and requests for resources and reagents should be directed to and will be fulfilled by the lead contacts, Nicholas A. Veldhuis, Daniel P. Poole and Karen J. Gregory

### Materials availability

The mGlu_5_-NP construct, HEK293A-mGlu_5-Venus_ and HEK293A-mGlu_5-RLuc8_ cell lines generated in this study are available from the lead contact upon request with a completed materials transfer agreement.

### Data and code availability

Any additional information required to reanalyze the data reported in this paper is available from the lead contact upon request.

## EXPERIMENTAL MODEL AND STUDY PARTICIPANT DETAILS

### Animals

Male C57BL/6 mice (6-10 weeks) were sourced from the Monash Animal Research Platform and the New York University Langone Medical Center. Male and female Sprague Dawley rats (8-12 weeks) were sourced from the Monash Animal Research Platform. Animals were maintained at 22 ± 3°C in a controlled environment of 12h light/dark cycle with food and water ad libitum. Studies were performed following the Guide for the Care and Use of Laboratory Animals of the National Institutes of Health and adhered to the ARRIVE guidelines.^63^ Investigators were double blinded to the treatment groups, and animals were randomly assigned to treatments and studied during the light cycle. At the end of the study, animals were euthanized by isoflurane overdose. The data presented in this study were derived from male mice and equal numbers of male and female rats. All animal care and experiments were approved by the Animal Ethics Committee of Monash Institute of Pharmaceutical Sciences, Monash University, or the Animal Ethics Committee of New York University Langone Medical Center.

### Cells

Monoclonal HEK293A cells stably expressing wild-type rat mGlu_5,_ at similar low levels to cortical astrocytes (hereafter referred to as HEK-mGlu_5_-low) have been previously described (Noetzel et al., 2013). Rat mGlu_5_ constructs containing C-terminal Venus (mGlu_5_-_venus_) or *Renilla* luciferase (mGlu_5_-_RLuc8_) were a gift from Kevin Pfleger (University of Western Australia). HEK293A cells (Invitrogen) were stably transfected with mGlu_5-Venus_ or mGlu_5-RLuc8_ using Lipofectamine 2000 transfection reagent (500 ng DNA, 1:4 DNA:Lipofectamine ratio) and selected using Dulbecco’s modified eagle medium (DMEM) with 5% (vol/vol) fetal bovine serum (FBS) and 500 μg/mL geneticin (G418). Single clones were selected and expanded, prior to confirmation of receptor and tag function. Cell lines described above, and non-transfected HEK293A cells, were maintained in DMEM with 5% FBS at 37°C and 5% CO_2_.

## METHOD DETAILS

### Constructs

Cytoplasmic EKAR and nuclear EKAR (Cerulean-Venus), hereafter referred to as cytoEKAR and nucEKAR, respectively, were gifts from Karel Svoboda (Addgene plasmid # 18679; http://n2t.net/addgene:18679; RRID:Addgene_18679: Addgene plasmid # 18681; http://n2t.net/addgene:18681; RRID:Addgene_18681). NES-Venus-mGα or NES-NLuc-mGα (mGα_s_, mGα_q/11_), HA-LgBiT-CAAX and HA-LgBiT-FYVE were kindly provided by Professor Nevin A. Lambert (Augusta University). mGlu_5_ with a C-terminal natural peptide tag (mGlu_5_-NP hereafter) was synthesized by Twist Bioscience (San Francisco, USA). mGlu_5_-NP consists of rat mGlu_5α_, followed by a linker (VPLGSSGGG) and NP tag (GVTGWRLCERILA)^64^. A similar approach was used to synthesize NK1R-NP, as a positive receptor control for endosomal signaling assays.

### Compounds

L-Glutamic acid, quisqualic acid, substance P and phorbol 12,13-dibutyrate (PDBu) were purchased from Merck (WI, USA). (S)-3,5-dihydroxyphenylglycine (DHPG), cyclopiazonic acid (CPA) and Dyngo4A were purchased from Abcam (Cambridge, UK). DL-threo-β-benzyloxyaspartic acid (DL-TBOA), JNJ16259685 and fenobam were purchased from HelloBio (Bristol, UK). DIPMA nanoparticles (DIPMA-NPs), which are polymeric nanocarriers responsive to acidic environments to deliver drugs were made in house as previously described^37^. VU0366058 was a gift from P. Jeffrey Conn (Vanderbilt University) and was synthesized as previously described^54^.

### Intracellular calcium mobilization assay (iCa^2+^)

HEK-mGlu_5_-low cells were seeded at a density of 30,000 cells/well into poly-D-lysine coated clear-bottom 96-well plates (Corning, NY, USA) using assay media (glutamine-free DMEM supplemented with 5% dialyzed FBS and 500 μg/mL G418) for 24 h. Cells were loaded with Fura2-AM ester (1 µM) in HEPES-buffered saline (150mM NaCl, 2.6mM KCl, 0.1mM CaCl_2_, 1.18mM MgCl_2_, 10mM D-glucose, 10mM HEPES and 0.5% BSA, pH 7.4) supplemented with 4 mM probenecid and pluronic F 127 (0.02%; Sigma Aldrich, Darmstadt, Germany) for 45 min at 37°C in a CO_2_-free incubator. Fura2-AM fluorescence was measured at 340 nm and 380 nm excitation and 530 nm emission, using a FlexStation I or III Microplate Reader (Molecular Devices, Sunnyvale, CA) as previously described^65–67^. A baseline of 20 s was recorded before agonist addition, with fluorescence recorded for a further 4 min. For inhibitor studies, cells were pre-incubated for 30 min with vehicle (0.3% DMSO), DL-TBOA (50 µM) or Dyngo4a (30 µM). For area under curve (AUC) analysis, raw kinetic fluorescence traces were plotted, and the total peak area in response to different concentrations of agonist and AUC from each experiment were normalized to responses to buffer (0%) and to ionomycin (1 μM; 100%).

### Cell transfection

For nuclear Ca^2+^, ERK1/2 FRET and BRET trafficking experiments, HEK-mGlu_5_-low or HEK-mGlu_5-RLuc8_ cells were seeded into a 10 cm Petri dish (Corning™, USA) at a density of ∼7×10^5^ cells in DMEM (with 5% FBS and 500 μg/mL G418) and incubated for 24 h (37°C, 5% CO_2_). Prior to transfection, media was changed to fresh DMEM (with 5% FBS). Cells were transfected with 2.5 μg DNA per dish using PEI at a 1:6 DNA:PEI ratio in 150 mM NaCl. Cells were then plated as outlined for each experiment.

### Nuclear Ca^2+^ responses

HEK-mGlu_5_-low cells transfected with the genetically encoded nuclear calcium indicator NLS-R-GECO a were seeded into poly-D-lysine coated clear 96-well culture plates (Perkin Elmer, USA) and incubated for 24h (37°C, 5 % CO_2_). Cells were equilibrated in Hanks balanced salt solution (HBSS; (MgCl_2_.6H_2_O 0.49 mM, MgSO_4_.7H_2_O 0.41 mM, KCl 5.33 mM, KH_2_PO_4_ 0.44 mM, NaCl 137.93 mM, Na_2_HPO_4_.7H_2_O (dibasic) 0.34 mM, D-glucose 5.56 mM) supplemented with 20 mM HEPES and 1.2 mM CaCl_2_, pH 7.4) at 37°C in a CO_2_-free incubator for 1 h. 10 U/ml glutamic pyruvic transaminase (GPT) and 6 mM sodium pyruvate were added for 1 h prior to assay to break down ambient glutamate. Cells were then pretreated for 30 min with vehicle (0.3% DMSO), DL-TBOA (50 µM), CPA (10 µM) or Dyngo4a (30 µM). NLS-R-GECO fluorescence was measured at 544nm excitation and 590nm emission using a FlexStation III Microplate Reader. Readings were captured every 1.5 s. A baseline of 20 s was recorded before agonist addition, followed by a 200 s read time post-addition. AUC was calculated from kinetic fluorescence plots. Data are reported as raw AUC.

### Förster resonance energy transfer (FRET)

HEK-mGlu_5_-low cells transfected with cytoEKAR or nucEKAR FRET biosensors were seeded into poly-D-lysine coated black 96-well culture plates (Perkin Elmer, USA) and incubated for 24h (37°C, 5 % CO_2_). Cells were serum-starved for 8-12 h prior to experiments. Cells were equilibrated in HBSS supplemented with 20 mM HEPES, 10 U/ml GPT and 6 mM sodium pyruvate at 37°C in a CO_2_-free incubator for 1 h. Transfected biosensors were excited at 430 nm and emission measured at 480 nm and 530 nm using a PHERAstar FS or FSX (BMG LABTECH, Germany). Measurements were recorded every 1 min, with a 5 min baseline period, followed by stimulation with agonist or positive control, PDBu (1 μM). Further measurements were taken for 30 min post-addition. FRET ratios (530 nm/480 nm) were calculated, baseline corrected and used to derive AUC.

### Bioluminescence resonance energy transfer (BRET)

HEK-mGlu_5-Rluc8_ cells transfected with miniGα_-Venus_, KRas_-Venus_ or RAB5_-Venus_ were seeded into poly-D-lysine coated white-wall, white-bottom 96-well plates and incubated for 24 h. On the day of the assay, cells were equilibrated in HBSS supplemented with 20 mM HEPES at 37°C in a CO_2_-free incubator. 10 U/ml GPT and 6 mM sodium pyruvate were added for 1 h prior to assay to break down ambient glutamate. Coelenterazine h (5 µM) was used as the substrate for RLuc8, added 10 min prior to recording. BRET was measured using a PHERAstar FS, with emission at 480 nm and 530 nm. Measurements were made every 1 min, with a 5 min baseline period, followed by stimulation with agonist. Further measurements were taken for 25 min post-addition. BRET ratios (530nm/480nm) were calculated, baseline corrected and used to derive AUC.

### NanoBRET studies of receptor signaling from cell surface and endosomal compartment

For NanoBiT-BRET (nbBRET) experiments investigating mGlu_5_ signal complex formation at the cell surface and in endosomes, HEK293A cells were seeded in 96-well white microplates at a density of 40,000 cells/well in DMEM with 5% FBS. After 4-6 h, cells were transfected with mGlu_5_-NP, LgBiT-CAAX or LgBiT-FYVE, and Venus-mGα_sq_ (10 ng per construct, 40 ng of total DNA per well, 1:6 DNA:PEI ratio in 150 mM NaCl) in a ratio of 1:1:2 or 1:2:1 for mGlu_5_-NP:LgBit-CAAX: Venus-mGα_sq_ and mGlu_5_-NP:LgBit-FYVE: Venus-mGα_sq_, respectively. Media was changed to assay media, 24 h post-transfection. After 48 h post transfection, cells were washed and pre-incubated with HBSS, supplemented with 20 mM HEPES, 10 U/ml GPT and 6 mM sodium pyruvate for 50 min, followed by 10 min incubation with NanoGlo® live cell substrate at a final concentration of 5µM. BRET was measured using a LUMIstar Omega or PHERAstar FS/FSX. Measurements were made every 1 min, with a 5 min baseline period, followed by stimulation with agonist and a further 25 min measurement period post-agonist addition. BRET ratios (530 nm/460 nm) were calculated and used to derive ΔBRET, the difference between the BRET signal after agonist and the baseline BRET signal. ΔΔBRET was further corrected for buffer response. AUC was calculated from ΔΔBRET.

### pH responsive nanoparticles (NPs)

Polymer synthesis, nanoparticle assembly, and characterization have been described previously.^37^ Briefly, self-assembling nanoparticles were made using a mixture of DIPMA polymer (10 µg/mL) and 1 mM VU0366058 in tetrahydrofuran (THF). The mixture was added into PBS (pH 7.4) under vigorous stirring. Finally, nanoparticles were dialyzed against PBS using dialysis membrane (MWCO 3500; ThermoFisher) for 24 h. DIPMA-Cy5 nanoparticles for imaging studies were assembled following the same methodology, without addition of drug, as described previously.^37^

### mGlu_5_ trafficking and immunofluorescence

HEK293-mGlu_5-Venus_ cells were seeded on poly-D-lysine coated glass coverslips in DMEM supplemented with 5% FBS for 24h before challenge with agonist (1 µM) for 5, 15, 30, or 60 min. Cells were washed once with PBS before fixation (4% paraformaldehyde, 15 min on ice), then incubated in permeabilization/blocking buffer (PBS containing 0.1% sodium azide, 0.1% saponin and 5% normal horse serum) for 1 h at room temperature. Cells were incubated with a monoclonal purified mouse anti-EEA1 (1:500; BD Biosciences) in blocking buffer overnight at 4°C, before washing and incubation with secondary antibody (donkey anti-mouse AlexaFluor 647, 1:500; ThermoFisher) in PBS for 2 h at room temperature. Coverslips were washed (3 x PBS), then mounted using ProLong Diamond Antifade Mountant (ThermoFisher) prior to imaging.

### Nanoparticle trafficking in HEK-mGlu_5_-low cells

HEK-mGlu_5_-low or HEK-mGlu_5-Venus_ cells were plated on poly-D-lysine coated glass coverslips in DMEM supplemented with 5% FBS for 24 h. Cells were incubated with DIPMA-Cy5 nanoparticles (20 μg/mL, 30 min, 37°C) or vehicle, followed by the addition of agonist (10 µM). Endosomes were identified by EEA1 immunofluorescence, as described above. The interaction of nanoparticles with endosomes after mGlu_5_ internalization was assessed in HEK-mGlu_5_-low cells, whereas HEK-mGlu_5-Venus_ was used to identify the interaction of nanoparticles with mGlu_5_ after agonist addition.

### Partial sciatic nerve ligation (PNL) model

A PNL was performed to injure the left sciatic nerve of 8-12 week old male and female Sprague-Dawley rats (n = 22, equal numbers of male and female) as described previously^68^. Briefly, rats were anaesthetized with isoflurane and the sciatic nerve proximal to its trifurcation was surgically exposed and a single suture was tied around one third to one half of the nerve. Rats were assessed for mechanical allodynia two weeks post-PNL surgery using a manual von Frey assay and used for electrophysiology experiments between 2-3.5 weeks post-surgery. A reduction in von Frey threshold from a pre-surgery baseline of ≥9.7g to ≤2.8 g 14 days after surgery was used as a threshold criterion that neuropathic pain had developed. Rats were housed in a temperature-controlled environment 22±2 °C with a 12 h light/dark cycle in groups of 3-4 and had free access to food and water.

### Spinal cord slice electrophysiology

Adult male and female Sprague-Dawley rats (10-16 weeks) that had undergone PNL were anaesthetized with isoflurane, decapitated, and the lumbar region of the spinal cord was removed. Parasagittal slices (340 μm thick) of spinal cord were cut on a vibratome in oxygenated ice-cold sucrose-based artificial CSF (saCSF; 100 mM sucrose, 63 mM NaCl, 2.5 mM KCl, 1.2 mM NaH_2_PO_4_, 1.2 mM MgCl_2_, 25 mM glucose, and 25 mM NaHCO_3_). Slices were transferred to a submerged chamber containing NMDG-based recovery ACSF (raCSF; 93 mM NMDG, 2.5 mM KCl, 1.2 mM NaH_2_PO_4_, 30 mM NaHCO_3_, 20 mM HEPES, 25 mM glucose, 5 mM sodium ascorbate, 2 mM thiourea, 3 mM sodium pyruvate, 10 mM MgSO_4_ and 0.5 mM CaCl_2_, pH 7.4), equilibrated with 95% O_2_ and 5% CO_2_ for 15 min at 34°C. Following recovery incubation, slices were transferred to normal oxygenated aCSF (125 mM NaCl, 2.5 mM KCl, 1.25 mM NaH_2_PO_4_, 1.2 mM MgCl_2_, 2.5 mM CaCl_2_, 25 mM glucose, and 11 mM NaHCO_3_) for 30 min at 34° C. Slices were then maintained at room temperature while equilibrated with 95% O_2_ and 5% CO_2_. Prior to recording, slices were incubated in DIPMA-VU0366058 (500 nM), VU0366058 (500 nM) or aCSF for 90-120 min at 34° C and then incubated in ACSF for 10 min. Some slices were pre-incubated with Dyngo4a (30 µM) for 10 min. Slices were transferred to a recording chamber and superfused continuously at 2 ml/min with oxygenated normal aCSF supplemented with NBQX (10 µM, Cayman), D-APV (50 µM, Cayman), gabazine (10 µM, SR95531, Abcam), strychnine (500 nM, Sigma) and the mGlu_1_ antagonist JNJ16259685 (100 nM). Perfusion aCSF was maintained at 34°C with an inline heater and monitored by a thermistor in the slice chamber. Dodt-contrast optics were used to identify laminae I and II neurons in the superficial dorsal horn. Neuron activity was recorded in whole-cell current clamp using a potassium gluconate-based internal solution (135 mM potassium gluconate, 10 mM HEPES, 0.5 mM EGTA, 8 mM NaCl, 2 mM MgATP, 0.3 mM NaGTP, 2 mM Lucifer Yellow CH dipotassium salt, 0.1% biocytin (osmolarity 285-295 mOsmol/L). Patch clamp electrodes had resistances of 6-9 MΩ. Input resistance was measured by injection of depolarizing current steps using a range that induced an output of -100 mV increasing in 10-40 pA increments until at least two steps above rheobase. Rheobase was measured by applying depolarizing pulses in 2-5mV increments until the neuron reached the action potential firing threshold. Following baseline recordings, either glutamate (100 µM) or DHPG (100 µM) were superfused onto cells and recordings repeated. Neurons were washed for 10-15 min in perfusion ACSF and recordings repeated. Neurons were filled with Lucifer Yellow and biocytin to allow identification of morphology at the end of the recording to ensure a similar range of neuronal subtypes were included in each treatment group.

#### Immunofluorescence in spinal cord

DIPMA-Cy5 nanoparticles were intrathecally administered as described above. Following nanoparticle administration (4 h), mice were transcardially perfused with 50mL of cold PBS followed by 50mL of 4% paraformaldehyde.^33,37^ The spinal cord was removed and fixed for 2 h at 4°C in 4% PFA and cryoprotected in PBS containing 30% sucrose (24 h at 4°C). The spinal cord (L3–L6) was embedded in tissue freezing medium (TFM, General Data), and 30 μm serial coronal sections were cut. Sections were washed twice in PBS and blocked in PBS containing 0.2% Triton X-100 and 10% normal horse serum (1 h, room temperature). Sections were incubated with mouse anti-HuC/HuD (1:500, ThermoFisher Scientific, A-21271) in PBS containing 0.2% Triton X-100 and 3% normal horse serum (overnight, 4°C). Sections were washed four times in PBS and incubated with donkey anti-mouse Alexa488 (1:1,000, ThermoFisher Scientific; 1 h, room temperature). Sections were counter-stained with DAPI (5μg/mL, 5 min) and mounted using ProLong Diamond mounting medium (ThermoFisher Scientific).

#### Confocal microscopy

Spinal cord sections or fixed cells were imaged using a super-resolution TCS-SP8 Lightning confocal microscope (Leica) equipped with HCX PL APO ×63 (NA 1.40) glycerol objective. For each treatment, three regions of interest were captured at 16-bit depth and 1024 × 1024- pixel resolution, with four independent experiments per treatment. Images were analyzed using Fiji, and co-localization of different markers evaluated by determining the Manders overlap coefficient.^69^

### *In vivo* pain assays

#### Drug administration

Male C57BL/6 mice (6-10 weeks) were used for all *in vivo* studies. Drugs used *in vivo* were fenobam (200 nM), VU0366058 (100-300 nM) or nanoparticles delivering an equivalent dose of VU0366058 (DIPMA-VU0366058, 20-60μg/mL, 100-300 nM) or vehicle (aCSF or 0.9% NaCl). All drugs were administered by intrathecal injection (5μl) into the intervertebral space (L4/L5) 30 min before intraplantar capsaicin (0.5 µg in 0.9% NaCl) administration, 48h after intraplantar complete Freund’s adjuvant (CFA, 0.5 mg/mL in 0.9% NaCl) administration or 10 days post sciatic nerve ligation for the acute inflammatory model, chronic inflammatory model and spared nerve injury model, respectively. 0.9% NaCl was used as vehicle control for intrathecal injections for both inflammatory models.

#### Acute inflammatory pain model

The acute nociceptive pain model was carried out as previously described.^33,37^ Briefly, 10μl capsaicin (5μg prepared in 80% v/v NaCl (0.9% m/v) and 20% v/v Tween20) or vehicle (0.9% NaCl) were subcutaneously administered in the plantar left hind paw of sedated mice (3% isoflurane). 30 min after capsaicin injection, drugs were administered intrathecally as described above.

#### Chronic inflammatory pain model

The inflammatory chronic pain model was carried out as previously described.^37^ Briefly, CFA (200µL of mycobacterium at 1 mg/mL) was mixed with an equal volume (200 µL) of saline solution (0.9% NaCl) to form an emulsion. CFA emulsion (10 µL) was subcutaneously administered in the plantar left hind paw of sedated mice (3% isoflurane), and after 48 h post CFA administration, drugs or nanoparticles were administered intrathecally as described above.

#### Spared nerve injury (SNI) Model

The SNI procedure was performed as previously described^70^, with modifications. Briefly, animals were anesthetized (5% inhaled isoflurane), the left thigh was shaved and the area sterilized using isopropyl alcohol and iodine wipes. A 1 cm skin incision was made on the lateral surface of the left thigh and the muscle bluntly dissected to identify the three terminal branches of the sciatic nerve. The common peroneal and tibial nerves were ligated using 6-0 silk (DemeTECH, USA), and a 1 mm segment of each nerve removed distal to the ligature. For sham procedures, the trifurcation was identified, but the nerves left untouched. The muscle layer was closed using 5-0 vicryl (Ethicon, USA), and the skin closed with a 5-0 Surgipro suture (Ethicon, USA). Buprenorphine (0.05 mg/kg, i.p.) was administered prior to cessation of anesthesia and then dosed every 12 h for 3 days post-surgery. Mice were randomly assigned to treatments or surgery and investigators were blinded to the surgery type (SNI or sham) and treatments (nanoparticles or free drug). All experiments were performed during the light cycle.

#### Mechanical allodynia

Mechanical nociception was assessed by measuring withdrawal thresholds of the ipsilateral and contralateral hind paw using calibrated Von Frey Filaments (VFF) as previously described.^33,37,46^ Briefly, before experiments commenced, animals were acclimatized for 2 h in individual acrylic boxes. A baseline of withdrawal thresholds was measured before drug or nanoparticle administration to establish a normal response for each mouse. For the acute inflammatory model, VFF withdrawal thresholds were measured at 30 min intervals for the first 3 h after drug administration, then at 60 min intervals for the next 2 h. For the chronic inflammatory model, VFF withdrawal thresholds were measured every 30 min for the first 3 h after drug administration, then at 60 min intervals for the next 3 h. For the neuropathic pain model, VFF withdrawal thresholds were measured every 30 min for 5 h after drug administration. Results are expressed as a percentage of baseline responses.

## QUANTIFICATION AND STATISTICAL ANALYSIS

Data were analyzed using GraphPad Prism 10 (GraphPad Software). Agonist concentration response curves from pharmacological assays in HEK293A cells were fitted to a three-parameter Hill equation:

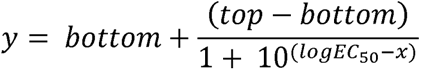

where ‘top’ and ‘bottom’ represent the maximum and minimum plateaus respectively, and ‘EC_50_’ represents the concentration of quisqualate required to evoke a 50% maximal response.

Data are presented as mean ± s.e.m., unless otherwise noted. For multiple comparisons, results were compared using one- or two- way ANOVA followed by post-hoc multiple comparison tests, as described in the figure legends. Differences were deemed statistically significant if *p* < 0.05.

## Results

### Glutamate induces mGlu_5_ internalization into early endosomes through dynamin-dependent pathways

As with many other GPCRs, mGlu_5_ undergoes endocytosis upon agonist activation through both clathrin- and caveolar/lipid raft mediated mechanisms^26,28,71^. However, the fate of internalized mGlu_5_ and the ability of mGlu_5_ to continue to signal from endocytotic compartments remains understudied. We first characterized the endocytosis profile of mGlu_5_ in response to the endogenous orthosteric agonist glutamate using bioluminescence resonance energy transfer (BRET). HEK293A cells stably expressing mGlu_5_ C-terminally tagged with *Renilla* luciferase protein 8 (HEK-mGlu_5-RLuc8_) were transiently transfected with GFP-tagged markers for the plasma membrane protein KRas (KRas_-GFP_) or the early endosomal membrane protein Rab5 (Rab5_-GFP_)^61^ (**Figure 1A**). Upon stimulation with glutamate, a rapid, sustained and concentration-dependent decrease in mGlu_5-RLuc8_ - KRas_-GFP_ BRET signal was observed, indicating mGlu_5_ dissociation from the plasma membrane (**Figure 1A, Supplementary Figure 1**). The decrease in mGlu_5_-KRas BRET correlated with a concentration-dependent, sustained increase in mGlu_5-RLuc8_-Rab5_-GFP_ BRET, indicating that mGlu_5_ internalizes into early endosomes (**Figure 1A, Supplementary Figure 1**).

**Figure 1.**
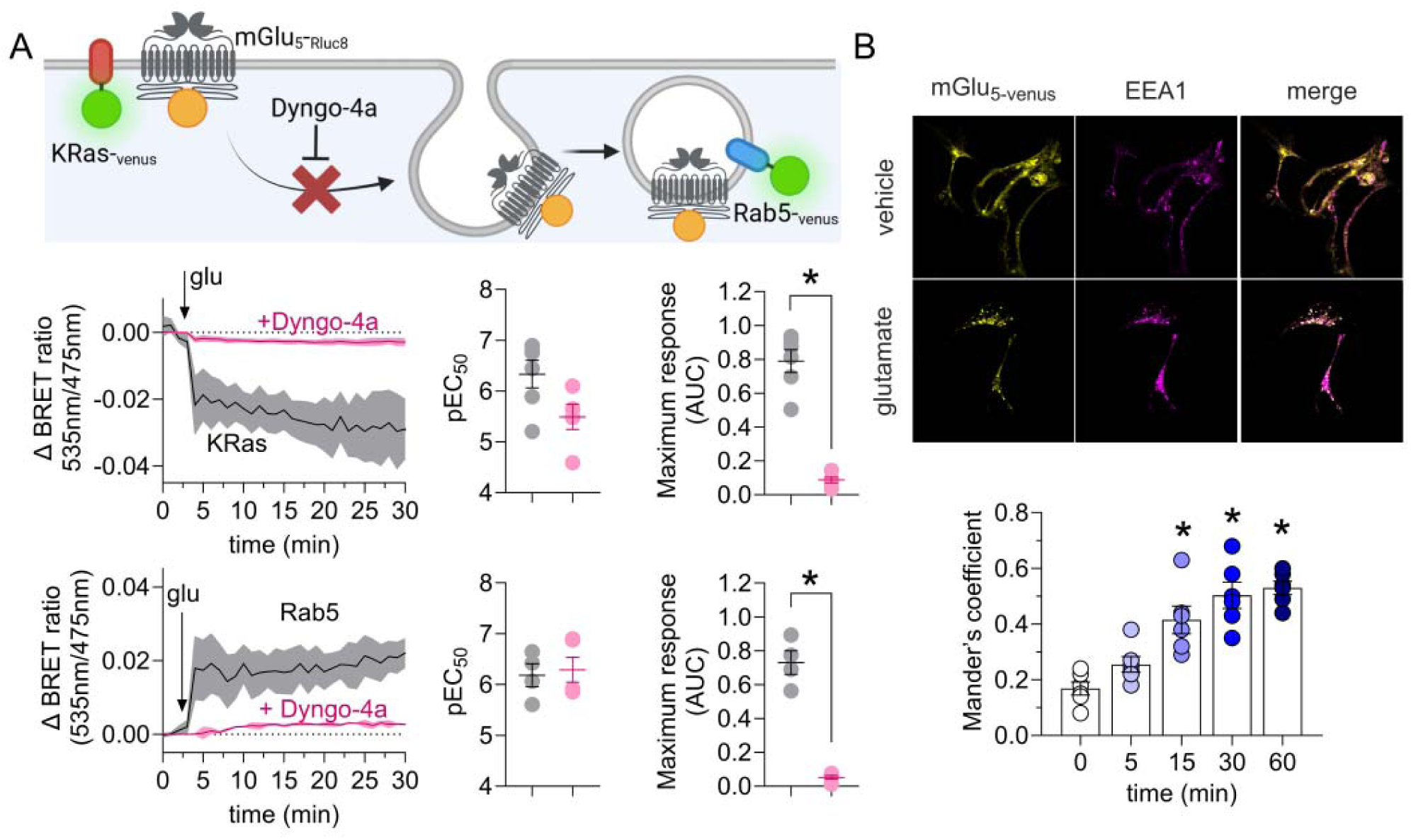
mGlu_5_ internalizes to early endosomes in a dynamin dependent manner. **A)** Kinetic traces of mGlu_5_ movement away from the cell surface and toward early endosomes, expressed as a change in BRET ratio for mGlu_5-RLuc8_ dissociation from KRas_-Venus_ or association with Rab5_-Venus_ upon glutamate (10µM) addition in absence (black) or presence (pink) of Dyngo-4a (30 µM). Scatter plots represent pEC_50_ and E_max_ values derived from glutamate concentration response curves shown in Supplementary Figure 1. Data are presented as mean ± s.e.m., n = 4-6 independent experiments performed in duplicate. \**p* < 0.05 compared to no Dyngo-4a, one-way ANOVA, Šídák’s multiple comparisons post-hoc test. **B)** Representative images of mGlu_5-Venus_ association with the early endosomal marker EEA1 upon vehicle or glutamate (1µM) treatment. Mander’s overlap coefficients show the time course of glutamate-induced mGlu_5_ movement towards early endosomes. Data are expressed as mean ± SEM of n = 6 independent experiments, 4-6 cells per experiment. \**p* < 0.05 compared to 0 min, one-way ANOVA, Dunn’s post-hoc test.

As both clathrin- and caveolar- mediated endocytosis involve the GTPase dynamin, we next probed the contribution of dynamin to mGlu_5_ endocytosis using the selective dynamin inhibitor Dyngo4A^73^. Pretreatment with Dyngo4A (30 µM) prevented glutamate-induced endocytosis, with a decrease in mGlu_5_ dissociation from the plasma membrane and association to early endosomes (**Figure 1A**). In the presence of Dyngo4A, glutamate E_max_ estimates were significantly decreased for mGlu_5_-Kras (**Figure 1A**).

We next confirmed early endosomal localization of internalized mGlu_5_ using confocal microscopy in HEK293A cells expressing mGlu_5_ with a C-terminal Venus tag (HEK293A-mGlu_5_-_Venus_). HEK293A-mGlu_5_-_Venus_ cells were labeled with an antibody for the early endosomal marker, early endosome antigen 1 (EEA1). Spectral overlap between Venus/EEA1 was quantified using Manders’ overlap coefficient analysis^69^ **(Figure 1B)**. In the absence of glutamate, mGlu_5-Venus_ fluorescence did not significantly overlap with EEA1 fluorescence (**Figure 1B**). However, in cells challenged with glutamate (1 µM), mGlu_5_ association with early endosomes increased over time, with a significant sustained increase in Manders’ overlap coefficient evident after 15 min (**Figure 1B**). Taken together, these data show that mGlu_5_ is rapidly endocytosed into early endosomes following agonist addition, in a dynamin dependent manner.

### mGlu_5_ recruits Gα_q_ and Gα_s_ with different spatiotemporal profiles

Consistent with prior spatiotemporal studies of GPCR signaling, we hypothesized that internalization of mGlu_5_ recruits G proteins to early endosomes to initiate divergent signaling events.^33,34,38,41,43,45^ While mGlu_5_ canonically couples to Gα_q/11_, there is also evidence to support coupling to Gα_s_ in recombinant systems^74,75^. Using BRET to confirm mGlu_5_ G-protein coupling profiles, we initially focused on global mGlu_5_ G protein recruitment, using stably expressing mGlu_5_-_Rluc8_ cells (HEK293-mGlu_5_-_Rluc8_) with transient expression of NES-Venus-mGα_s_ or NES-Venus-mGα_q/11_^59^ (**Figure 2A**). Stimulation with glutamate resulted in a rapid, concentration-dependent increase in BRET between mGlu_5_-_Rluc8_ and Venus-tagged mGα_s_ and mGα_q/11_ proteins and elevated BRET was maintained for the duration of the measurement period (28 min post-stimulation, **Figure 2A, Supplementary Figure 2**). Notably, glutamate-induced recruitment of mGα_q/11_ was biphasic (**Figure 2A**). Concentration response curves revealed equal glutamate potency for mGα_s_ or mGα_q/11_ protein recruitment to mGlu_5_, whereas the E_max_ for mGα_q/11_ was reduced due to the biphasic nature of the response. (**Figure 2A**). We next investigated mGlu_5_-dependent recruitment of G proteins to endosomes. In stable untagged HEK-mGlu_5_ cells transiently co-expressing mGα_s_- or mGα_q/11_-NES - RLuc8 with RAB5_-venus_ as the BRET acceptor, glutamate addition increased mGα_q/11_-RLuc8/RAB5-_Venus_ BRET signals, but not mGα_s_-RLuc8/RAB5-_Venus_ (**Figure 2B**). Confocal microscopy was used to confirm the recruitment of mini-G proteins into early endosomes seen in BRET studies. HEKA cells with stable expression wild type mGlu_5_ and transient expression of mGα_s_ or mGα_q/11_-_Venus_ were stained with an anti-EEA1 antibody to detect early endosomes. Consistent with BRET data, the distribution of mGα_q/11_, but not mGα_s_, co-distributed with EEA1-positive structures after glutamate addition (**Figure 2C**). Together, BRET and microscopy studies are consistent with glutamate-induced mGlu_5_ internalization and recruitment of mGα_q/11_ protein into the endosomal network.

**Figure 2.**
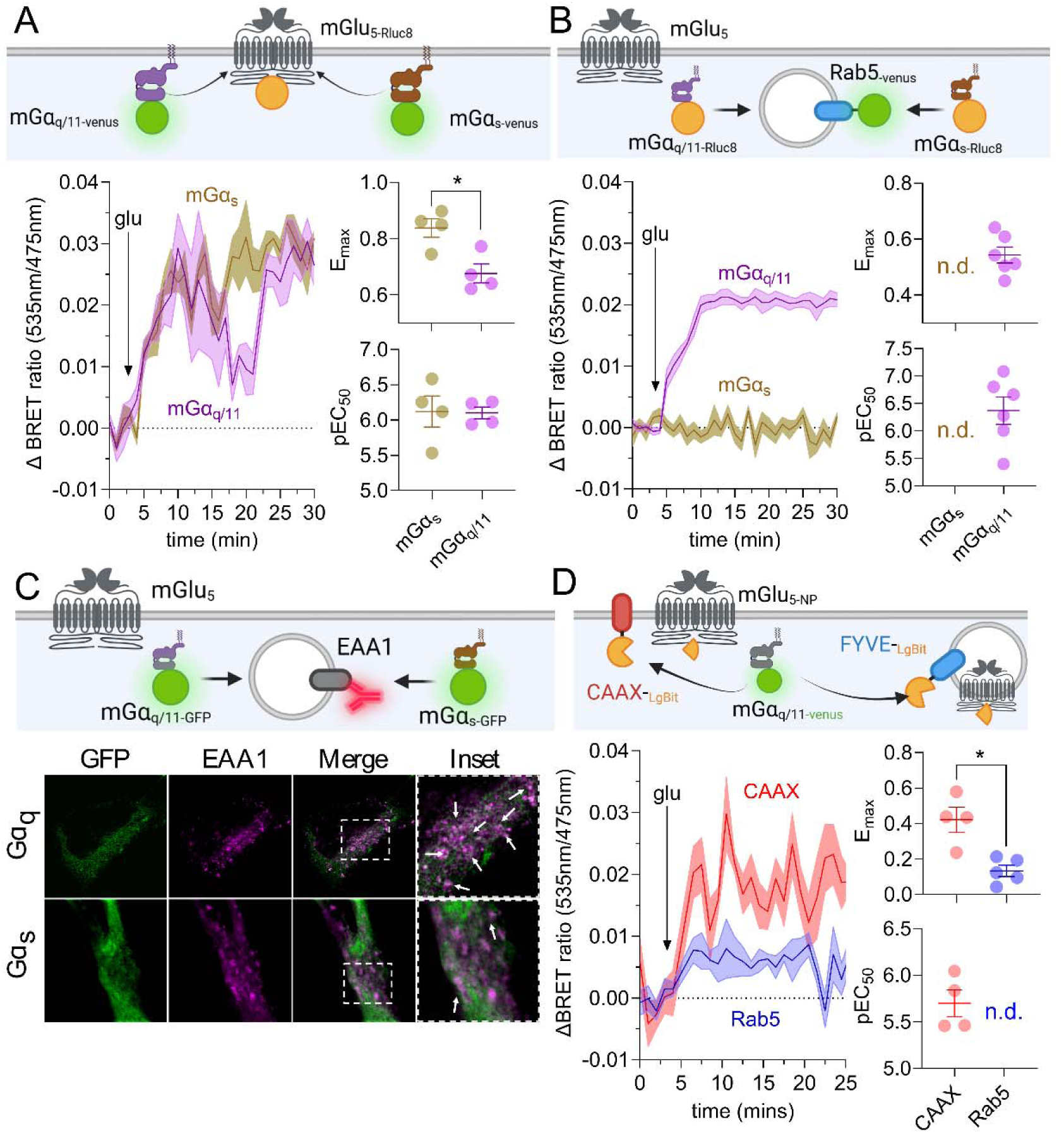
mGlu_5_ recruits mini-Gα_q/11_ to both the cell surface and endosomes upon activation. **A)** Kinetic traces of G protein recruitment to mGlu_5_, expressed as change in BRET ratio for mGlu_5-RLuc8_ association with mG_αs-Venus_ (brown) or mG_αq/11-Venus_ (purple) upon glutamate (100 µM) addition. **B**) Kinetic traces of G protein recruitment to endosomes, expressed as change in BRET ratio for Rab5_-Venus_ association with mG_αs-RLuc8_ (brown) or mG_αq/11-RLuc8_ (purple) upon glutamate (100 µM) addition. Scatter plots represent pEC_50_ and E_max_ values derived from glutamate concentration response curves (see Supplementary Figure 2). **C)** Representative confocal microscopy images of mG_αs-GFP_ or mG_αq/11-GFP_ recruitment to endosomes (green). Endosomes are labeled with an antibody for the endosomal marker EEA1 (purple). **D)** Kinetic traces of G protein recruitment to mGlu_5_ on the cell surface or in early endosomes, expressed as a change in BRET ratio for mG_αq/11-Venus_ association with mGlu_5-NP_ + CAAX-LgBit (cell surface, red) or mGlu_5-NP_ + FYVE-LgBit (early endosome, blue) upon glutamate (100µM) addition. Data are presented as mean ± s.e.m., n = 4-6 independent experiments performed in duplicate. \**p* < 0.05, one-way ANOVA, Šídák’s multiple comparisons post-hoc test. N.D. not determined due to lack of response.

The BRET and imaging experiments above show that mGlu_5_ activity elevates G protein proximity to Rab5 or EEA1-enriched membranes. However, this does not measure direct recruitment of mGα_q/11_-_Venus_ to mGlu_5_-positive endosomes. To address this limitation, we used the nanobit-BRET (nbBRET) complementation system to measure mGlu_5_ recruitment of mGα_q/11_ specifically to the cell surface or early endosomes^64^ (**Figure 2D**). mGlu_5_ was C-terminally tagged with the natural peptide NLuc sequence (mGlu_5_-_NP_) and co-transfected with LgBiT fused to targeting sequences for the plasma membrane (CAAX-LgBiT) or early endosome (FYVE-LgBiT). When co-expressed with Venus-mG_αq/11_ as the NLuc acceptor, this enables exclusive nbBRET-based measurement of mGlu_5_-dependent G protein recruitment to the cell surface or early endosomes. As a positive control, we tested a GPCR with known endosomal coupling and signaling, the neurokinin 1 receptor (NK_1_R), which was also tagged with NP (NK_1_R-_NP_). The NK_1_R agonist Substance P (SP) induced a concentration dependent increase in BRET between NK_1_R-_NP_ / CAAX-LgBiT and mGα_q/11-Venus_ and between NK_1_R-_NP_ / FYVE-LgBiT and mGα_q/11-Venus_, confirming validity of the BRET constructs and approach (**Supplementary Figure 3**). For mGlu_5_-_NP_, stimulation with glutamate increased nbBRET at the plasma membrane (**Figure 2D, Supplementary Figure 2**). In contrast, only the highest concentration of glutamate (100 µM) increased nbBRET between mGlu_5_-_NP_, FYVE-LgBiT and mGα_q/11-Venus_ (**Figure 2D, Supplementary Figure 2**). While these results from nbBRET experiments confirmed our observations of endosomal G protein recruitment from global BRET and microscopy studies, these data indicate that a relatively minor population of mGlu_5_ is able to directly recruit mG_αq/11_ to endosomes.

### Sustained mGlu_5_ signaling requires receptor internalization

mGlu_5_ stimulation activates distinct signaling events in discrete subcellular locations, such as the nucleus^20,22,25,47^. Furthermore, excitatory amino acid transporter 3 (EAAT3) actively transports glutamate into the cell from the extracellular space, affecting signaling by intracellular mGlu_5_ ^7,14,21,76,77^. However, little is known about the exact contribution of mGlu_5_ internalization to these signaling events. We next characterized location-specific mGlu_5_ iCa^2+^ and extracellular regulated kinase 1/2 (ERK1/2) signaling profiles to determine the contribution of internalization to global and nuclear mGlu_5_ signaling, using fluorescent calcium dyes and genetically encoded localized sensors, respectively. Assays were carried out in the absence or presence of dynamin inhibitors to prevent mGlu_5_ internalization. While Dyngo4A was used for iCa^2+^ mobilization, Dyngo4A fluorescence can interfere with FRET-based assays, therefore expression of a dominant negative dynamin K44E was used to inhibit internalization for ERK1/2 FRET assays. Mutations to dynamin residue K44 effectively block early steps of the endocytotic pathway and inhibit receptor endocytosis ^78^. As HEK293A cells endogenously express EAAT3^79^, the EAAT3 inhibitor DL-TBOA was used to investigate how glutamate transport into the cell affects mGlu_5_ signaling.

Stimulation of cells with glutamate triggered global iCa^2+^ mobilization consisting of a rapid first phase (0 – 30 s post-addition) and a second, sustained phase (>30 s post-addition) **(Figure 3B**). The second phase was characterized by significantly lower glutamate potency and E_max_ estimates compared to the first phase **(Figure 3, Supplementary Figure 4)**. Inhibition of mGlu_5_ internalization with Dyngo4A (30µM, 30 min) decreased E_max_ of both phases of glutamate-induced iCa^2+^ mobilization compared to vehicle treatment response, without affecting glutamate potency (**Figure 3, Supplementary Figure 4**). We next assessed cytosolic ERK1/2 activity using the cytoEKAR FRET biosensor. Stimulation with glutamate induced a rapid and sustained biphasic increase in cytosolic ERK1/2 activity, measured using the cytoEKAR FRET biosensor (**Figure 3C, Supplementary figure 4**). The two phases were defined as rapid (0-10 min post-addition, first phase) and sustained (>10 min post-addition, second phase). Inhibition of dynamin-mediated internalization by co-transfection with mutant K44E slowed the rate of the rapid phase of ERK1/2 activity and decreased E_max_ estimates for the sustained, second phase of cytosolic ERK activity (**Figure 3C & E**). Taken together, iCa^2+^ mobilization and ERK1/2 activation assays revealed the importance of mGlu_5_ internalization for sustained global mGlu_5_ signaling.

**Figure 3.**
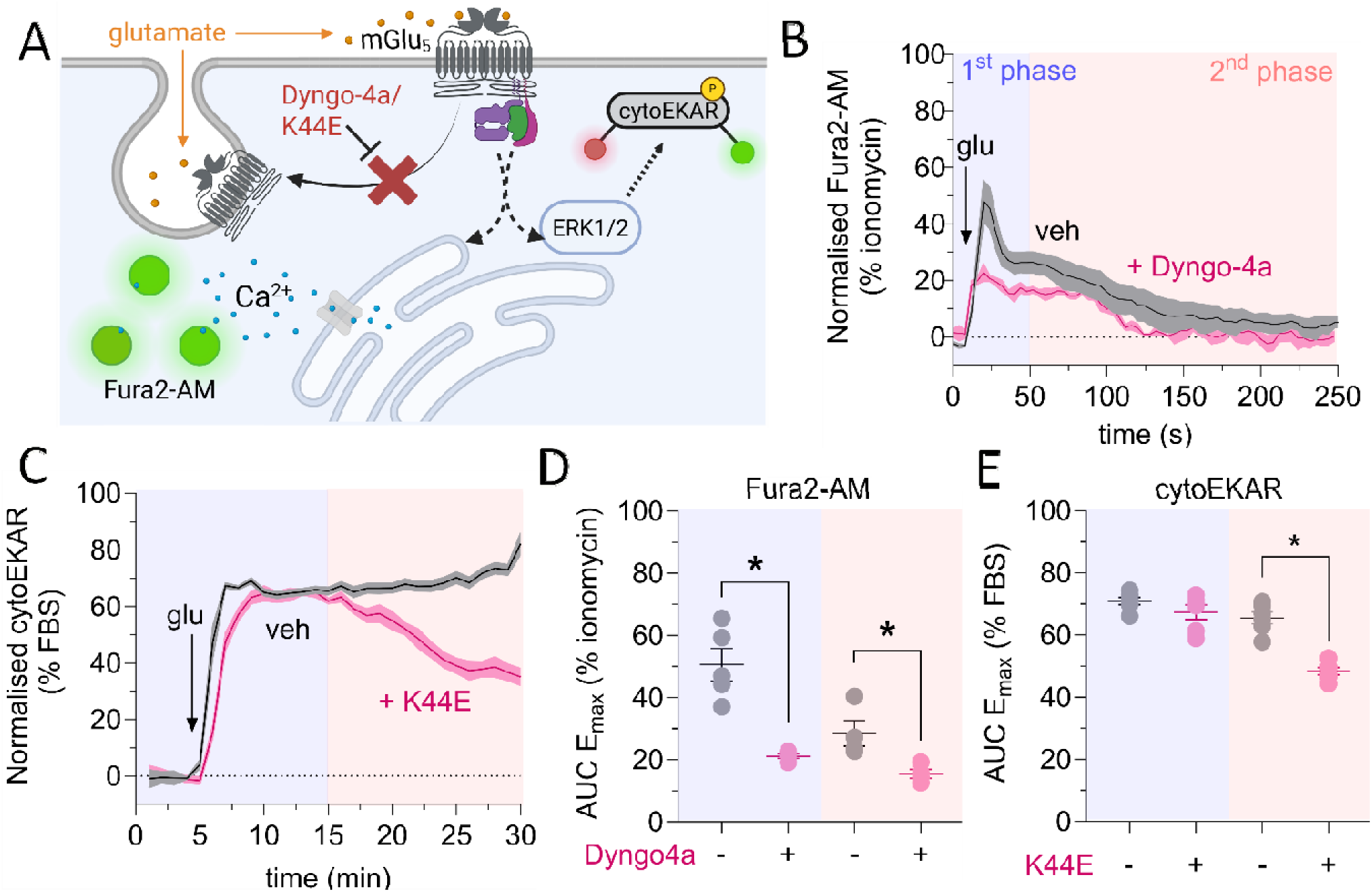
Sustained global mGlu_5_ signaling relies on receptor internalization in HEK293A cells. **A)** Schematic of tools and sensors used in global signaling assays. **B-C)** Kinetic traces of glutamate (10µM) evoked **B**) global intracellular Ca^2+^ mobilization, measured using Fura2-AM, and **C**) cytosolic ERK1/2 activity, measured using the cytoEKAR biosensor. Cells were preincubated with vehicle (0.3% DMSO, black), Dyngo-4A or K44E (30µM, pink). **D-E)** Scatter plots representing E_max_ values derived from glutamate concentration response curves shown in Supplementary Figure 4 divided into first phase (blue background) and second phase (red background). \**p* < 0.05, one-way ANOVA, Šídák’s multiple comparisons post-hoc test. N.D. not determined due to lack of response. Data are presented as mean ± s.e.m., n = 4-6 independent experiments performed in duplicate.

### Glutamate-induced nuclear mGlu_5_ iCa^2+^ and ERK1/2 signaling are differentially regulated by receptor internalization and glutamate transport

We next studied the link between mGlu_5_ endocytosis and localized signaling from the nucleus. mGlu_5_ is associated with activation of nuclear signaling through both iCa^2+^ mobilization and ERK1/2 pathways. However, it remains unknown how mGlu_5_ internalization affects these processes^7,14,22,23^. We assessed glutamate-induced nuclear Ca^2+^ mobilization and ERK1/2 activation using a nuclear localization sequence-tagged genetically-encoded Ca^2+^ indicator (NLS-R-GECO) and nucEKAR biosensors, respectively^56,80^. Dynamin-mediated endocytosis and glutamate transport were inhibited as described above.

Glutamate triggered a biphasic Ca^2+^ mobilization response, characterized by a rapid increase over the first 30 s post-addition, followed by a decrease before a second, sustained phase from >30 s post-addition (**Figure 4B, Supplementary Figure 4**). As such, responses were divided into first (acute, 0 – 30 s post-addition) and second (>30 s post-addition) phases for analysis. Given the rapidity of the initial phase, we hypothesized that nuclear Ca^2+^ flux is linked to cell surface receptor activation via the endoplasmic reticulum (ER), as ER membranes express mGlu_5_ and are contiguous with nuclear membranes. ER Ca^2+^ stores were depleted using cyclopiazonic acid (CPA, 30 µM) which inhibits the Sarcoplasmic/Endoplasmic Reticulum Ca² -ATPase (SERCA) pump, a transporter of Ca^2+^ from the cytosol into the endoplasmic reticulum^81^. Preincubation with CPA significantly inhibited the maximum response of both phases of the nuclear Ca^2+^ response, without affecting glutamate potency **(Figure 4B, Supplementary Figure 4C**), indicating that ER-derived Ca^2+^ plays a key role in nuclear responses to mGlu_5_ stimulation. Inhibition of mGlu_5_ internalization with Dyngo4A significantly reduced the E_max_ for the second phase of the nuclear Ca^2+^ response without affecting the first phase, with no effect on glutamate potency for either phase (**Figure 4B & D, Supplementary Figure 4**). Glutamate transport is not required for nuclear Ca^2+^ mobilization, as EAAT3 inhibition with DL-TBOA did not significantly affect glutamate potency or E_max_ for either phase (**Figure 4B, Supplementary Figure 4C**). Combining DL-TBOA and Dyngo4A did not significantly affect either phase when compared to Dyngo4A alone (**Figure 4B, Supplementary Figure 4**).

**Figure 4.**
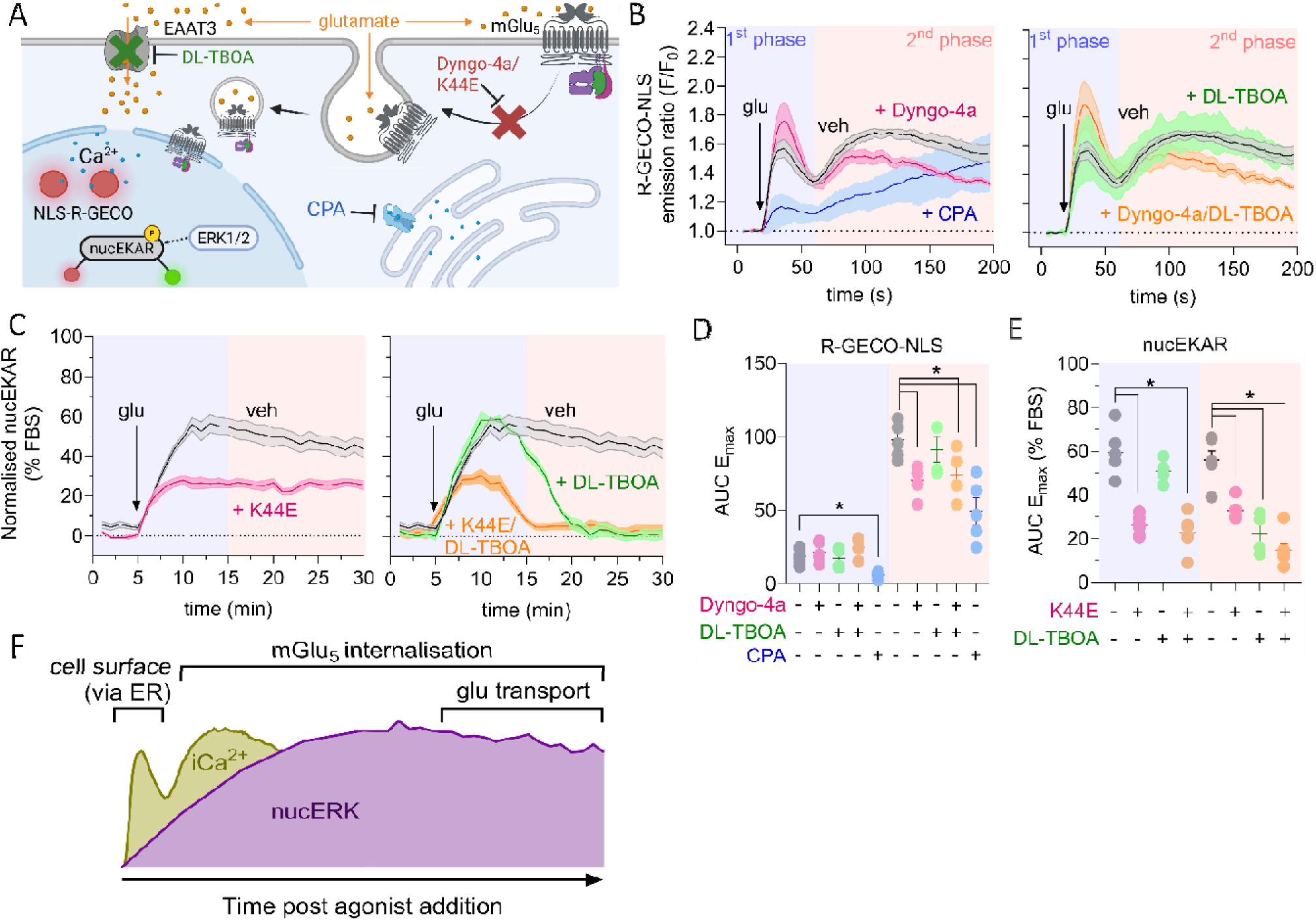
Sustained nuclear mGlu_5_ signaling relies on receptor internalization and glutamate transport in HEK293A cells. **A)** Schematic of tools and sensors used in nuclear signaling assays. **B-C)** Kinetic traces of glutamate (10µM) evoked **B**) nuclear intracellular Ca^2+^ mobilization, measured using R-GECO-NLS, and **C**) nuclear ERK1/2 activity, measured using the nucEKAR biosensor Cells were preincubated with vehicle (0.3% DMSO, black), Dyngo-4A or K44E (30µM, pink), CPA (30µM, blue), DL-TBOA (50µM, green) or both Dyngo-4A/K44E and DL-T OA (orange). **D-E)** Scatter plots representing E_max_ values derived from glutamate concentration response curves shown in Supplementary Figure 4 divided into first phase (blue background) and second phase (red background). **F)** schematic representation of nuclear mGlu_5_ Ca^2+^ and ERK1/2 and the contribution of mGlu_5_ internalization and glutamate transport \**p* < 0.05, one-way ANOVA, Šídák’s multiple comparisons post-hoc test. N.D. not determined due to lack of response. Data are presented as mean ± s.e.m., n = 4-6 independent experiments performed in duplicate.

Nuclear ERK1/2 activation was also characterized by a sustained response, peaking 5 min after glutamate-stimulation and remaining elevated for the entire 25 min assay period **(Figure 4C)**. Responses were again divided into first (acute, 0-10 min post-addition) and second (sustained, >10 min post-addition) phases for analysis. Inhibition of mGlu_5_ internalization with K44E mutant dynamin resulted in a blunted first phase, but sustained signaling was still evident, albeit with lower E_max_ (**Figure 4C**). Glutamate potency and E_max_ estimates for both phases were significantly reduced in the presence of K44E (**Figure 4E, Supplementary Figure 4**). In contrast, glutamate transport was only required for sustained signaling. Preincubation with DL-TBOA did not impact the first phase response compared to vehicle, but nuclear ERK1/2 signaling was not sustained, returning to baseline within 15 min post-glutamate addition (**Figure 4C**). DL-TBOA significantly reduced glutamate potency and E_max_ for the second phase, without significantly affecting first phase responses (**Figure 4C & E, Supplementary Figure 4**). When combined with K44E, DL-TBOA did not affect first phase nuclear ERK1/2 activation when compared to K44E alone, but had a synergistic effect on inhibition of the second phase, with abolition of nuclear ERK1/2 activation within 10 min post-glutamate addition (**Figure 4C & E, Supplementary Figure 4**).

Together, these data indicate that glutamate-induced intracellular signaling by mGlu_5_ is dependent on either mGlu_5_ endocytosis, glutamate transport into the cytosol, or both, depending on the signaling pathway measured. Importantly, Ca^2+^ mobilization is acute (signaling over ∼3 min) when compared to extended ERK1/2 activation (∼25 min). Examination of these two signaling measures allows us to construct a timeline of nuclear mGlu_5_ signaling following glutamate stimulation (**Figure 4F**). Glutamate activation of nuclear mGlu_5_ causes a rapid Ca^2+^ peak (0-1 min post-stimulation), which is driven through the ER and does not require mGlu_5_ endocytosis or glutamate transport. The sustained nuclear Ca^2+^ response overlaps with the first phase of nuclear ERK1/2 signaling (2–5 min post-stimulation), both of which are mediated in part by mGlu_5_ internalization but not glutamate transport. Sustained nuclear ERK1/2 signaling (10-25 min post-stimulation) requires both mGlu_5_ internalization and glutamate transport into the cytosol. Sustained signaling also aligns with the kinetics of mGlu_5_ trafficking to endosomes (**Figure 1**), implicating mGlu_5_ localization, trafficking and endosomal signaling in sustained signaling responses.

### Nanoparticles containing an mGlu_5_ NAM associate with endosomal compartments

Nanoparticles are gaining interest as a means to selectively release drug-like molecules into specific intracellular locations, such as endosomes. Modulation of GPCR signaling in endosomes using NP-delivery of drugs decreases pain behavior ^36,37,41,45,82^. mGlu_5_ internalizes into early endosomes, where it can couple to Gα_q/11_ to contribute to sustained signaling, as demonstrated by our *in vitro* data. We developed dimethylaminoethyl methacrylate (DIPMA) nanoparticles loaded with a negative allosteric modulator (NAM) of mGlu_5_ as a tool for selective targeting of endosomal mGlu_5._ DIPMA nanoparticles are pH-responsive, rapidly releasing their contents in acidic environments such as those in endosomes, thereby allowing subcellular targeting of drug action^37^

DIPMA nanoparticles were self-assembled from block copolymers consisting of a hydrophilic (P(PEGMA-co-DMAEMA) and hydrophobic component p(DIPMA-co-DEGMA). Drug loaded pH-responsive nanoparticles were co-assembled in aqueous solution with copolymers and VU0366058 (VU058), a hydrophobic NAM of mGlu_5_.^54^. Referred to hereafter as DIPMA-VU058, transmission electron microscopy images indicated DIPMA-VU058 nanoparticles were present as a uniform spherical shape (**Figure 5A**) and dynamic light scattering (DLS) quantified a mean diameter of 44.4 ± 1.5nm with a slight negative ζ potential (-0.41 ± 0.05mV). Liquid chromatography-mass spectrometry (LC-MS) analysis indicated a loading efficiency of 40% (**Figure 5B**). Empty DIPMA particles were a similar spherical shape, with a smaller diameter of 38.6 ± 3.4nm and comparable ζ potentials (-0.5 ± 0.02mV).

**Figure 5.**
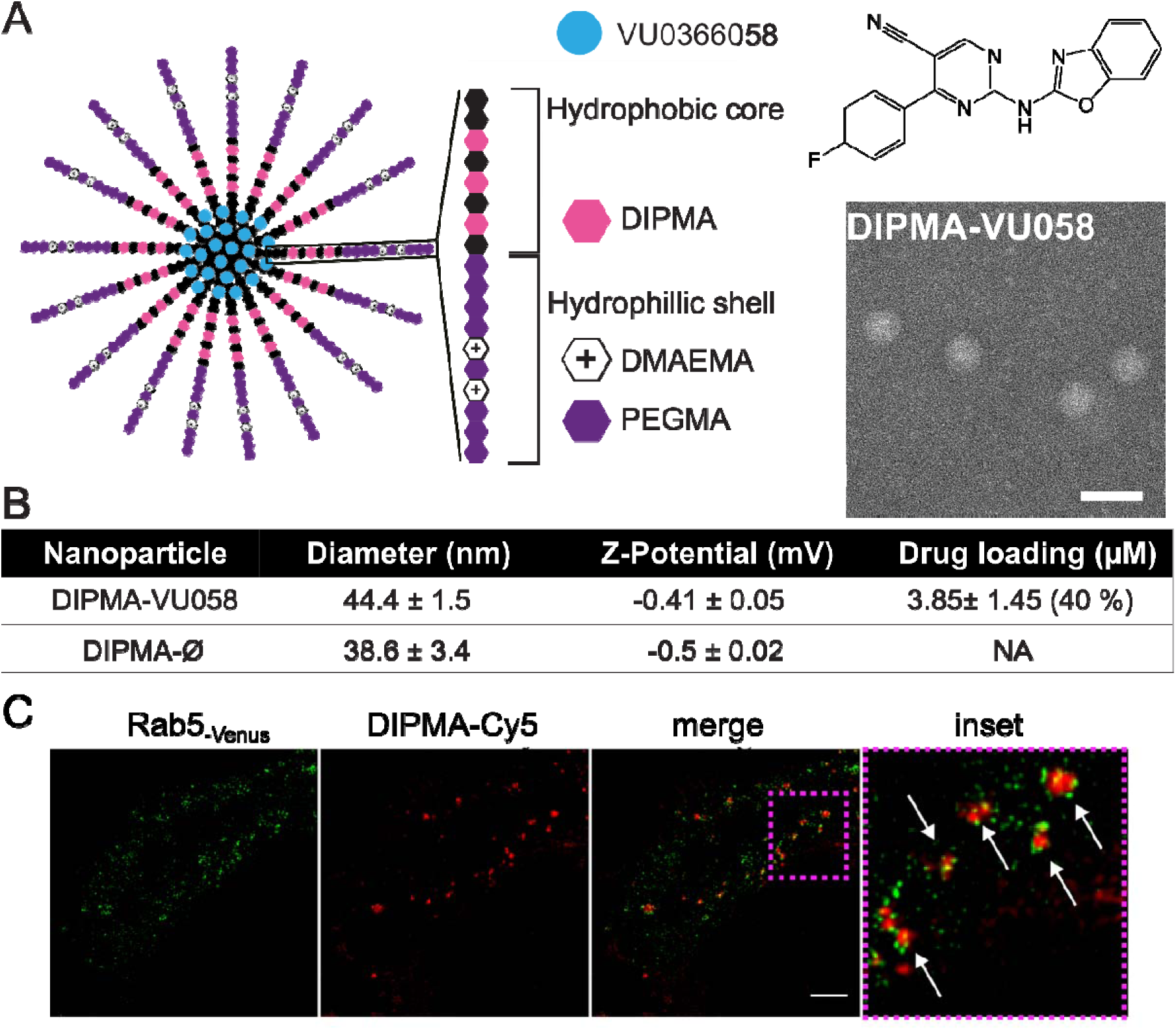
Characterization of DIPMA nanoparticles. **A)** Structure of pH responsive (DIPMA) nanoparticles, comprised of a hydrophilic shell of P(PEGMA-co-DMAEMA) and the hydrophobic cores of P(DIPMA-co-DEG A). The chemical structure of the mGlu_5_ negative allosteric modulator VU0366058 (VU058) is also shown. Representative transmission electron microscopy image shows DIPMA-VU058 (DIPMA loaded with VU0366058) nanoparticles. **B)** Properties of DIPMA-VU058 and DIPMA-Ø, determined by dynamic light scattering and LC-MS. Values in parenthesis indicate the percentage of the initial drug incorporated into DIPMA particles (% initial VU0366058). Data are presented as mean ± SD of n = 9 experiments. NA, not applicable. **C)** Representative confocal microscopy images of HEK293A cells transfected with Rab5_-GFP_ and incubated with DIPMA-Cy5 nanoparticles for 30 min.

To confirm the capacity for these nanocarriers to access the endosomal network, the cellular fate of DIPMA particles in HEK293A cells was examined by confocal microscopy. DIPMA nanoparticles labeled with the far-red-fluorescent dye cyanine 5 (DIPMA-Cy5) were incubated with HEK293A cells transiently expressing Rab5_-GFP_, a resident protein of the early endosome. DIPMA-Cy5 nanoparticles significantly co-distributed with Rab5+ endosomes 15 min post-DIPMA addition, as previously reported^37^ (**Figure 5C**). With prior studies demonstrating that equivalent DIPMA-based micelles offer rapid pH-responsive cargo release, these results confirm that DIPMA particles can readily internalize into cells to facilitate drug targeting within the endosomal network.

### An intracellular targeted mGlu_5_ NAM enhances inhibition of sustained mGlu_5_ signaling

Internalized mGlu_5_ contributes to sustained Ca^2+^ influx and nuclear ERK activity. We next examined whether nanoparticle encapsulated VU0366058 inhibits global and localized mGlu_5_ signalling. First, to confirm that DIPMA nanoparticles had potential to accumulate within mGlu_5_-specific endosomes, mGlu_5-Venus_ cells were pre-incubated Cy5-labelled DIPMA nanoparticles for 15 min, before challenging cells with vehicle or glutamate. When cells were challenged with glutamate, DIPMA-Cy5 particles significantly overlapped with mGlu5-_Venus_ when examined at 20 min post-stimulation (**Figure 6A & B**). We next assessed the capacity for DIPMA-VU058 to modulate mGlu_5_ endosomal signaling, using Ca^2+^ and nuclear ERK assays respectively. Any modulation of signaling can be attributed to VU0366058 as we have previously demonstrated that empty DIPMA (DIPMA-Ø) particles do not activate, or effect agonist-activated, nuclear ERK in HEK293A cells.^37^

**Figure 6.**
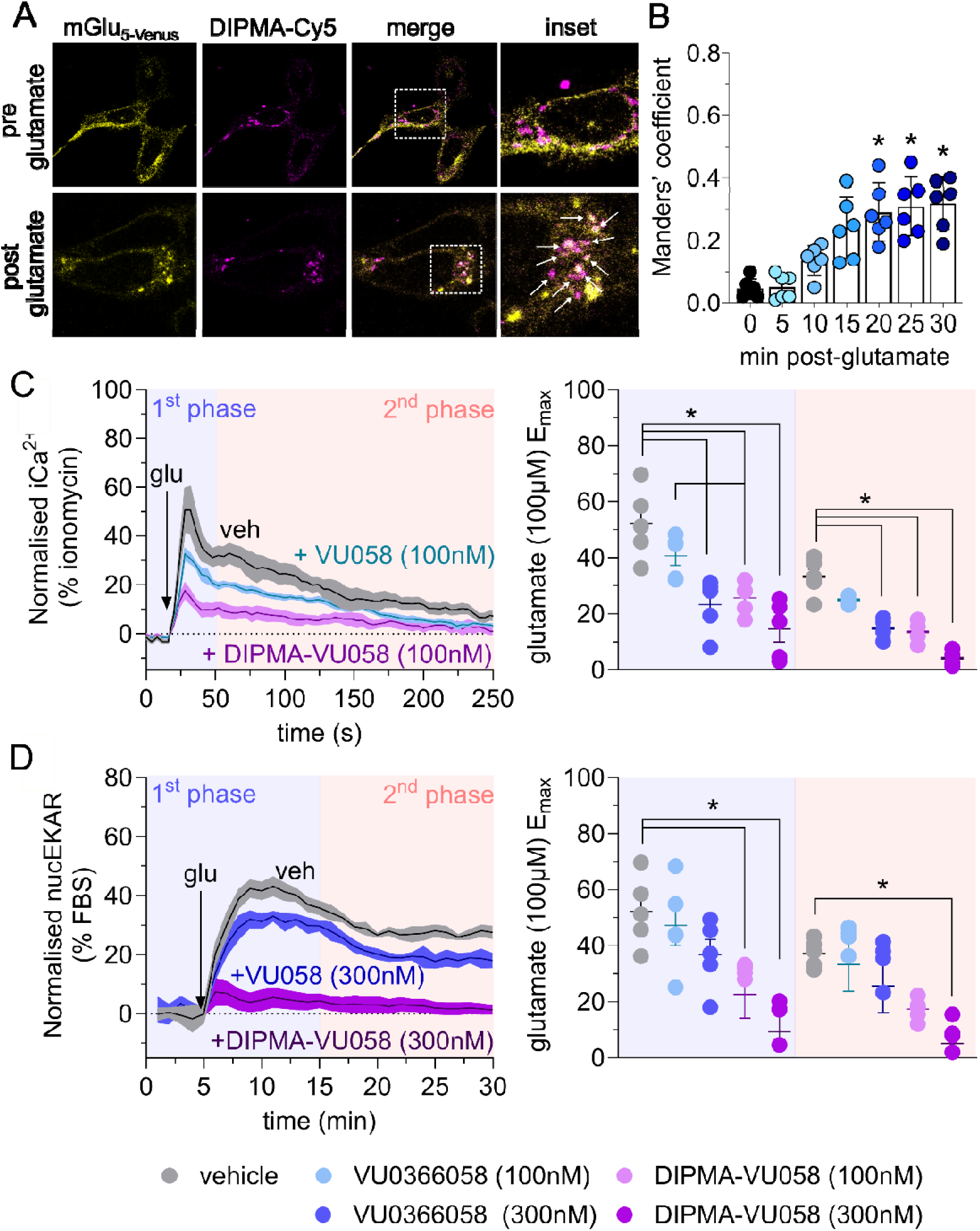
Nanoparticle encapsulation enhances VU0366058 inhibition of glutamate-induced cytosolic iCa^2+^ and nuclear ERK1/2 signaling in HEK293A cells. **A)** Representative confocal microscopy images of HEK293A cells transfected with mGlu_5-venus_. Cells were incubated with DIPMA-Cy5 nanoparticles for 15 min, then challenged for 30 min with vehicle or 1 µM glutamate to promote mGlu_5_ internalization. **B**) Manders’ overlap coefficients for degree of colocalization of DIPMA-Cy5 with mGlu_5-venus_. **C)** Kinetic traces of iCa^2+^ mobilization, measured using Fura2 dye, upon glutamate (100µM) addition in the presence of vehicle (black), free VU0366058 (100nM, blue) or DIPMA-VU058 (100nM, purple). **D**) Kinetic traces of nuclear ERK1/2 activity, measured using the nucEKAR biosensor, upon glutamate (100µM) addition, in the presence of vehicle (black), free VU0366058 (300nM, blue) or DIPMA-VU058 (300nM, purple). Scatter plots represent E_max_ values derived from AUC analysis of kinetic traces for 100µM glutamate for each condition, divided into first phase (blue background) and second phase (red background). Imaging data are presented as mean ± s.e.m., n = 6 independent experiments. *p < 0.05, **p < 0.005, ***p < 0.001, ****p < 0.0001 compared to 0 mins, one-way ANOVA, Dunn’s post-hoc test. Signaling data presented as mean ± s.e.m., n = 4-6 independent experiments performed in duplicate. *p < 0.05, one-way ANOVA, Šídák’s multiple comparisons post-hoc test.

HEK293A-mGlu_5_ cells were preincubated with vehicle, VU0366058 or DIPMA-VU058 (100 or 300 nM) for 30 min and stimulated with increasing concentrations of glutamate. In vehicle treated cells, glutamate induced a rapid increase in Ca^2+^ mobilization followed by a sustained phase, 30 – 250 s post-stimulation (**Figure 6B, Supplementary Figure 5**). Free VU0366058 had distinct, concentration-dependent effects on each phase of Ca^2+^ mobilization. Pre-treatment with 300 nM of free or DIPMA-formulated VU0366058 significantly decreased glutamate-induced Ca^2+^ mobilization in both signaling phases when compared to vehicle. In contrast, at 100nM, only DIPMA-VU058 significantly inhibited the first Ca^2+^ mobilization phase (**Figure 6B**). Group data comparing the E_max_ showed that inhibition with 100 nM of DIPMA-VU058 was comparable to 300 nM free VU0366058 in both phases of the Ca^2+^ signaling.

In HEKA cells stably expressing mGlu_5_ with transient expression of nucEKAR, glutamate stimulated a rapid and sustained activation of nuclear ERK1/2. Similar to iCa^2+^ assays, free VU0366058 at submaximal concentrations resulted in only partial inhibition of nuclear ERK1/2 activation, with little effect on the sustained second phase (**Figure 6C**). AUC analysis of kinetic traces revealed no significant effect of free VU0366058 on the glutamate E_max_ (100µM) for either phase compared to vehicle (**Figure 6C, Supplementary Figure 5**). DIPMA-VU058 inhibited glutamate-induced nuclear ERK1/2 activation to a greater extent than free VU0366058 at both 100nM and 300nM, with DIPMA-VU058 (300nM) again abolishing the second phase (**Figure 6C**). AUC analysis revealed significant decreases in glutamate E_max_ in the first phase for both concentrations of DIPMA-VU058, with only a significant effect of 300nM DIPMA-VU058 in the second phase (**Supplementary Figure 5**). Together, nanoparticle characterization and functional assays suggest nanoparticle encapsulation successfully targets VU0366058 to internalized and endosomal mGlu_5_, effectively inhibiting endosomal signaling and modulating both global and localized mGlu_5_ signaling cascades.

### Nanoparticle delivery of VU0366058 reduces glutamate and DHPG-induced excitability of dorsal horn neurons

To investigate the effect of mGlu_5_ activation on neuronal excitability of nociceptive neurons, we first performed electrophysiological recordings from neurons within the superficial laminae of spinal cord slices, taken from rat models of neuropathic pain. To isolate mGlu_5_ responses, we superfused slices with ionotropic glutamate receptor antagonists (D-APV 10 µM and NBQX 50 µM, to prevent AMPA, kainate and NMDA receptor activation), inhibitory receptor antagonists (strychnine 10 µM, gabazine 500nM) and an mGlu_1_ antagonist (JNJ16259685 100nM). As no differences were seen between neurons from male or female rats (equal numbers of animals used), data are grouped in the following figures and analyses. Following baseline recordings to measure resting membrane potential, neuronal input resistance and rheobase, glutamate (100 µM) was bath applied, which significantly depolarized membrane potential when assessed at 2 min post-application **(Figure 7A)**. Resting membrane potential returned to baseline levels 10-15 min following washout. Significant increases in neuronal excitability were evident by the decrease in the current required to elicit an action potential (rheobase) from 45.7 ± 5.7 pA to 17.14 ± 10.1 pA (*p* = 0.008, **Figure 7A**). Similar neuronal responses were observed 2 min after bath application of the mGlu_1/5_ selective agonist DHPG (100 µM) with significant, but reversible changes in resting membrane potential (7.25 mV depolarization, *p* = 0.007), input resistance (increased by 103 MΩ, *p* = 0.024) and rheobase (decreased by 24.6 pA, *p* = 0.029) **(Figure 7B)**. This excitation of spinal neurons is consistent with the sustained Ca^2+^ response and nuclear ERK1/2 signaling mediated by internalized mGlu_5_.

**Figure 7.**
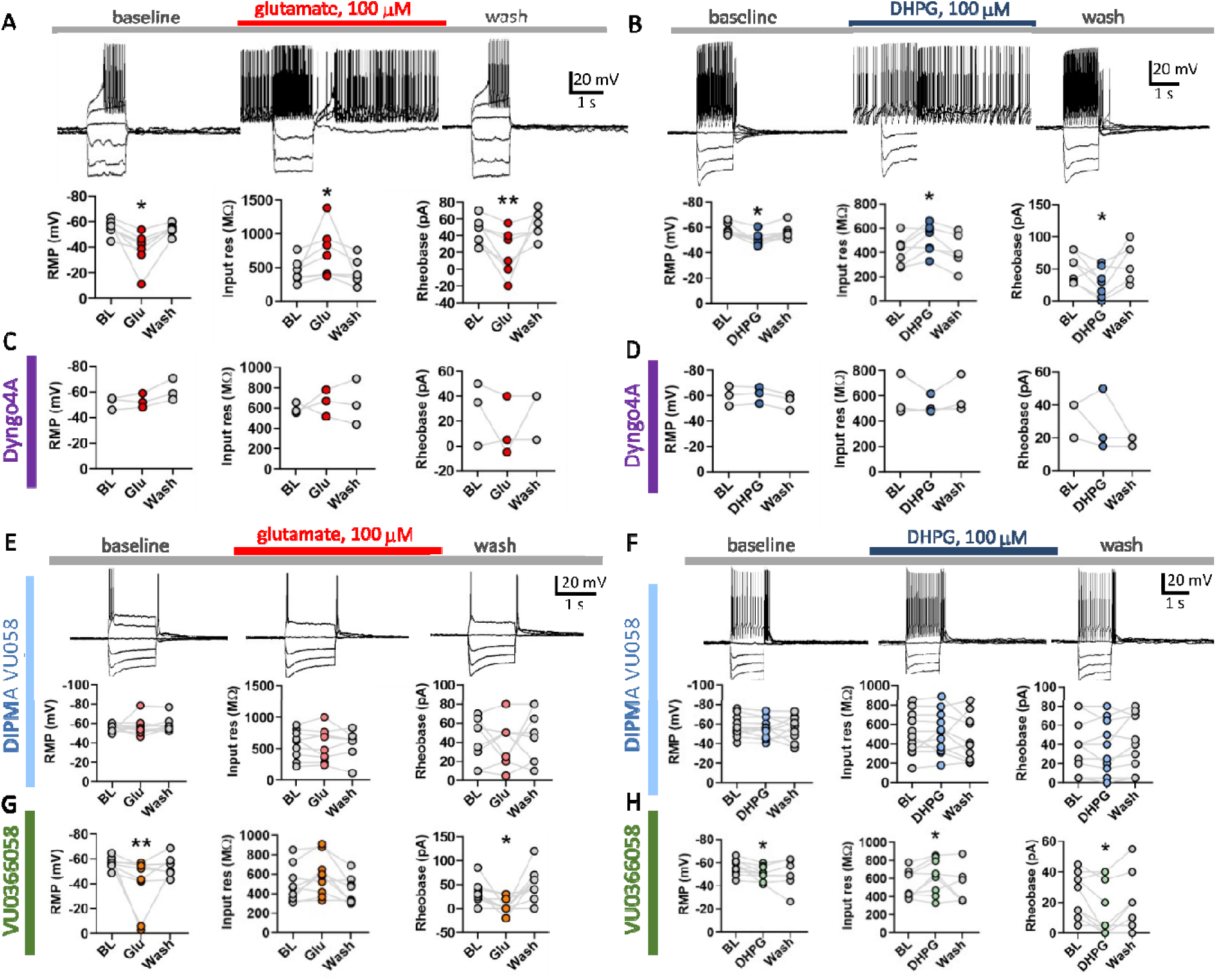
VU0366058 encapsulated in nanoparticles prevents mGlu_5_-agonist mediated hyperexcitability in spinal neurons from rats with neuropathic pain. **A-B)** Representative whole cell patch-clamp recordings from neurons in the superficial laminae of the lumbar spinal cord show mGlu_5_-mediated excitation in response to **A**) 100 µM glutamate (n = 7 neurons) or **B**) 100 µM DHPG (n = 7). Resting membrane potential (RMP), input resistance and rheobase were recorded at baseline (BL) prior to agonist superfusion, 2 min post glutamate or DHPG, and again 10-15 min following washout (wash). **C-D) F**ollowing 10 min pre-incubation in Dyngo4a (30µM) baseline activity was recorded before exposure to **C**) glutamate (n = 3) or **D**) DHPG (n = 3), followed by washout. Dyngo4a prevented glutamate and DHPG-induced excitability. **E-F)** Following a 90-min pre-incubation with DIPMA-VU058 (500 nM), baseline activity was recorded before exposure to **E**) glutamate (n = 9) or **F**) DHPG (n = 10), followed by washout. DIPMA-VU058 prevented glutamate and DHPG-induced excitability **G-H)** Slices were pre-incubated with free VU0366058 (500 nM) prior to recording and subsequent exposure to **G**) glutamate (n = 9) or **H**) DHPG (n = 10). Free VU0366058 failed to affect glutamate and DHPG-induced changes in excitability. All recordings were performed in the presence of synaptic receptor antagonists (NBQX, D-APV, strychnine and gabazine) and mGlu_1_ antagonist (JNJ16259685). No differences were seen between neurons from male or female rats (equal numbers of animals used). \**p* < 0.05, \*\**p* < 0.005, agonist compared to baseline, paired t test for those with normal distribution, or Wilcoxon matched-pairs signed rank test for those without normal distribution, normality was determined using a Shapiro-Wilk test.

First, to investigate the importance of mGlu_5_ endocytosis on excitability, slices were pre-treated with the dynamin inhibitor Dyngo4A (30 µM) for 10 min to prevent endocytosis. Dyngo4A prevented the effects of glutamate and DHPG on membrane potential, input resistance and rheobase (**Figure 7C & D**). These data confirm that endocytosis and intracellular pools of mGlu_5_ are critical for enhancing the excitability of spinal cord neurons. To assess if endosomal mGlu_5_ specifically contributes to sustained excitability and can be selectively modulated using nanoparticle delivery, slices were pre-incubated for 90 min with DIPMA-VU058 or free VU0366058 (500 nM) prior to recording. Following incubation with DIPMA-VU058, there was no significant difference in resting membrane potential, input resistance or rheobase when slices were superfused with glutamate (100 µM, **Figure 7E**) or DHPG (100 µM, **Figure 7F**). In contrast, changes in excitability were still observed in slices incubated with a molar equivalent of VU0366058, with significant depolarization induced by glutamate (16.92 mV, p = 0.007, **Figure 7G**) and a small but significant increase with DHPG (4.6 mV, p = 0.03, **Figure 7H**), a significant increase in input resistance with DHPG (93.6 MΩ, p = 0.04) and significant decreases in rheobase for both glutamate and DHPG (26.6 pA, p = 0.02; 9.5 pA, p = 0.03, respectively). This is consistent with Dyngo4A inhibition of neuronal excitability and also HEK293A signaling data, where DIPMA-VU058 significantly enhanced inhibition of mGlu_5_-dependent global Ca^2+^ mobilization and nuclear ERK activity compared to free VU036605 in recombinant cells. These data show nanoparticle-encapsulated VU0366058 can successfully target intracellular mGlu_5_ to reduce neuronal excitability in nociceptive circuits.

### Nanoparticle delivery of VU0366058 enhances analgesic efficacy in multiple nociceptive pain models

Based on our *in vitro* and electrophysiology data, we hypothesized that nanoparticle encapsulation would enhance the antinociceptive actions of VU0366058 by targeting endosomal mGlu_5_ in spinal neurons. As such, we next evaluated the efficacy of free or nanoparticle encapsulated VU0366058 in preclinical models of acute, inflammatory and neuropathic pain (**Figure 8**). Fenobam was used as a reference mGlu_5_ NAM with known antinociceptive properties^7,48,50,53,83,84^. Intrathecal injections (5 µL) were given to result in approximate local molar concentrations of each treatment; vehicle (0.9% w/v NaCl), DIPMA-VU058 (100 – 300 nM), free VU0366058 (100 – 300 nM) or fenobam (200 nM). Treatments were injected either before intraplantar capsaicin injection (acute model), after intraplantar complete Freund’s adjuvant injection (CFA, inflammatory model) or after sciatic nerve injury (neuropathic model). Mechanical nociception was studied by measuring withdrawal responses to stimulation of the plantar surface of the hindpaw using von Frey filaments. Mechanical withdrawal thresholds were defined as the minimum gauge Von Frey filament that elicited a withdrawal reflex. VFF threshold reduction is indicative of enhanced nociception. Given the lack of difference between mGlu_5_-mediated effects in male and female rats in electrophysiology studies, male mice were used for *in vivo* studies.

**Figure 3.**
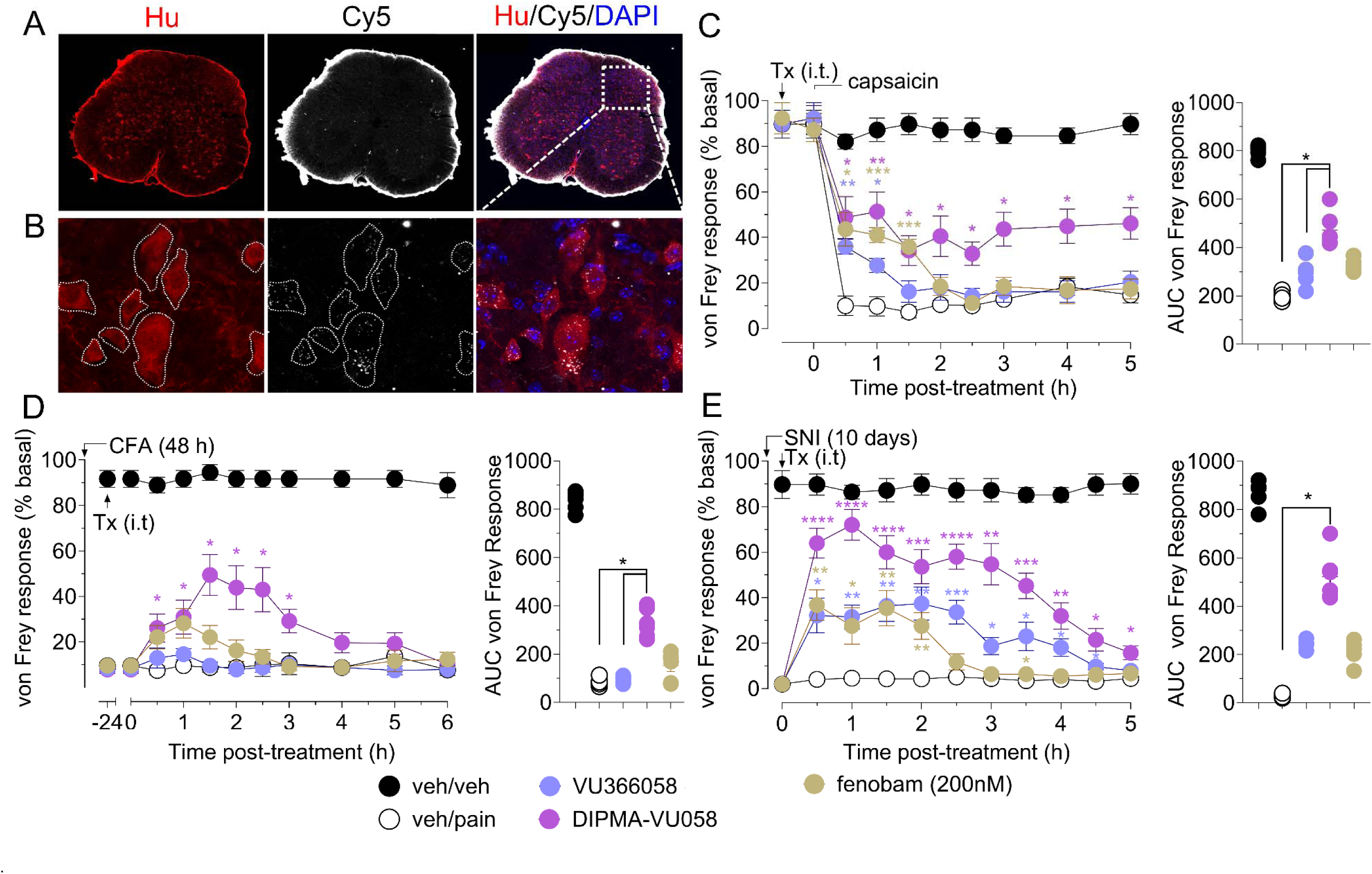
Nanoparticles internalize into spinal cord neurons and encapsulation enhances VU0366058-mediated analgesia in acute, inflammatory and neuropathic pain models in male mice. **A)** Confocal microscopy (40x objective) of spinal cord sections 4 h post intrathecal injection of Cy5-loaded DIPMA nanoparticles. Neurons are stained with mouse anti-HuC/HuD (red) and nuclei are stained with DAPI. **B)** 63x objective images of spinal cord neurons from the area marked with dashed square in panel A. Cy5 nanoparticles localize to punctate areas within spinal cord neurons, consistent with endosomal delivery. **C)** Free VU0366058 (100 nM) and fenobam (200 nM) resulted in modest reduction in capsaicin-evoked acute nociceptive pain, whereas DIPMA-VU058 (100nM) resulted in long-lasting analgesia over 5 h **D)** Neither free VU0366058 (300 nM) or fenobam (200nM) had any effect on Complete Freund’s adjuvant (CFA)-evoked sustained inflammatory pain, whereas DIPMA-VU058 (300nM) resulted in significant analgesia from 0.5-3 h post-intrathecal injection. **E)** DIPMA-VU058 (300 nM) treatment results in analgesia of a higher magnitude and duration than free VU0366058 (300 nM) or fenobam (200 nM) 10 days post SNI. Scatter plots represent integrated AUC responses for each pain model. Data are presented as mean ± s.e.m., n = 6 animals for all experiments. For time courses, \**p* < 0.05, \*\**p* < 0.005, \*\*\**p* < 0.001, \*\*\*\**p* < 0.0001 compared to veh/pain, two-way ANOVA, Dunnett’s multiple comparison post-hoc test. For scatter plots, \**p* < 0.05, one-way ANOVA, Kruskal-Wallis multiple comparisons post-hoc test.

To test efficiency of *in vivo* DIPMA-mediated delivery to spinal cord neurons, confocal microscopy of spinal cord sections was undertaken on tissues harvested 4 h post-intrathecal injection of Cy5-loaded DIPMA nanoparticles. Co-staining for HuC/HuD+ neurons revealed Cy5 signals localized to punctate areas within spinal cord neurons, consistent with endosomal delivery via DIPMA nanoparticles (**Figure 8A & B**).

Intraplantar capsaicin-induced hyperalgesia is mediated through mGlu_5_ via activation of transient receptor potential vanilloid 1 (TRPV1) on primary sensory neurons and subsequent glutamate release^85–88^. In mice pre-treated intrathecally with vehicle, capsaicin decreased the VFF threshold when assessed from 0.5 to 5 h post-capsaicin (**Figure 8C**). Pretreatment with fenobam (200 nM) caused a modest antinociceptive effect after 0.5 h (41 ± 14% inhibition), which was sustained for 1.5 h. Free VU0366058 (100 nM) had a similar effect after 0.5 h (35 ± 5% inhibition) which was sustained for 1.5h. However, DIPMA-VU058 (100 nM) had a marked antinociceptive effect from 0.5 h (54 ± 11% inhibition) that was sustained for 5 h (46 ± 7% inhibition). AUC analysis revealed that only DIPMA-VU058 (100 nM) resulted in significant reversal of acute pain over the entire time course when compared to vehicle. DIPMA-VU058 was also significantly more efficacious at inducing antinociception than free VU0366058 (**Figure 8C**)

Intraplantar injection of CFA causes inflammation-mediated sustained mechanical allodynia. There was a marked decrease in von Frey responses when assessed 48 h after intraplantar CFA injection (**Figure 8D**). Intrathecal administration of vehicle (0.9% NaCl) did not affect mechanical hyperalgesia, which persisted for 24 h. Fenobam administration (200 nM) transiently decreased mechanical hyperalgesia induced by CFA from 0.5 - 1.5 h, but this effect was not significantly different to vehicle. Free VU0366058 (300 nM) did not reverse hyperalgesia at any time. In contrast, DIPMA-VU058 (300 nM) induced a potent transient inhibition of mechanical hyperalgesia, significantly different to vehicle, starting from 0.5 h and maintained for 3 h. As in the acute model, DIPMA-VU058 induced antinociception was significantly different to both vehicle and free VU0366058 (**Figure 8D**)

The spared nerve injury (SNI) model produces a chronic mechanical hyperalgesia which persists for >50 days^89^. At 10 days post-surgery, SNI reduced the pressure withdrawal of the ipsilateral hindpaw when compared to sham-operated mice, indicative of mechanical hyperalgesia (**Figure 8E**). Intrathecal vehicle administration did not affect mechanical hyperalgesia. Fenobam (200 nM) administration significantly inhibited withdrawal thresholds after 0.5 h, to a maximum of 36 ± 8% inhibition, before returning to baseline after 2 h. Free VU0366058 (300 nM) also significantly reversed hyperalgesia after 0.5 h (32 ± 5% inhibition), with hyperalgesia returning to baseline after 3.5 h. DIPMA-VU058 (300 nM) resulted in a marked reduction of hyperalgesia, reaching a peak after 1 h (72 ± 6% inhibition), with significant analgesic effects maintained for 5 h (**Figure 8E**).

The enhanced effects of DIPMA-VU058 compared to free drug in multiple preclinical pain models is likely related to intracellular delivery and retention of VU0366058 in spinal neurons and the continued release of drug as nanoparticles encounter increasingly acidified endosomal compartments^37^. Targeting intracellular mGlu_5_ using nanoparticles therefore represents a viable strategy for enhancing the analgesic properties of mGlu_5_ NAMS to treat multiple types of pain.

## Discussion

Growing evidence indicates that GPCR signaling is not confined to the plasma membrane. GPCRs that traffic to endosomes or are natively present on organelles such as the nucleus or endoplasmic reticulum are signaling competent, and activating different receptor pools leads to distinct cellular signalling and functional outcomes ^7,14,33–37,90,91^. This is true for GPCRs involved in pain, including the neurokinin 1 receptor (NK1R), calcitonin receptor-like receptor (CLR), delta opioid receptor (DOR) and protease-activated receptor-2 (PAR2), for which endosomal receptor pools generate sustained signaling in primary sensory and spinal neurons to mediate nociception ^33–37,41,42,46,58^. mGlu_5_, a Class C GPCR, is also implicated in nociception and presents a new therapeutic target for the treatment of pain ^92^. While mGlu_5_ is known to signal from intracellular membranes of the nucleus and endoplasmic reticulum, the endosomal trafficking and signaling profile of mGlu_5_ and its contribution to pain pathology remains unknown. Here we report mGlu_5_ internalizes into early endosomes, where a small but significant pool couples to Gα_q_, but not Gα_s_, G proteins. Inhibition of mGlu_5_ internalization and active glutamate transport into the cytosol differentially affect cytosolic and nuclear Ca^2+^ and ERK1/2 signaling, with pain-related sustained nuclear signaling mediated by both processes. pH responsive nanoparticles were localized to endosomes in both recombinant cells and spinal cord neurons. Nanoparticle delivery of the mGlu_5_ NAM VU0366058 enhanced inhibition of cytosolic and nuclear mGlu_5_ signaling, reduced spinal cord neuron excitability, and provided more efficacious and longer lasting analgesia in multiple pain models. Our study is the first to demonstrate endosomal mGlu_5_ signaling and provides compelling evidence that targeting endosomal and intracellular mGlu_5_ with nanoparticle-encapsulated NAMs is an effective approach to treating pain.

mGlu_5_ internalizes both constitutively and upon agonist stimulation^26,93–96^. Various cellular mediators are involved in mGlu_5_ internalization and subsequent trafficking, including β-arrestin, Norbin, protein kinase C (PKC), G protein-coupled receptor kinases (GRKs), tamalin and calmodulin ^30,31,93,94,97–99^. Once internalized, mGlu_5_ traffics to early endosomes, before moving to recycling endosomes and returning back to the cell surface ^31,32,99,100^. Additionally, mGlu_5_ is expressed on various intracellular membranes, including those of the endoplasmic reticulum and nucleus, with up to 90% of mGlu_5_ localized intracellularly in neurons ^7,20,21,25,47,77,90^. Here, we demonstrate for the first time that mGlu_5_ continues to signal from early endosomes, with a small receptor pool able to recruit Gα_q_ after internalization. Canonically, β-arrestin is involved in GPCR internalization following activation by agonists, and plays an important role in sustained endosomal GPCR signaling ^101–103^. However, while β-arrestin is involved with constitutive mGlu_5_ internalization, agonist induced mGlu_5_ internalization is β-arrestin-independent ^29,104^. mGlu_5_ thus appears to belong to a growing group of GPCRs that challenge convention and do not require β-arrestin to mediate sustained endosomal signaling following internalization ^38,105–107^. Blocking mGlu_5_ internalization with Dyngo-4a also reduced nuclear signaling, indicating that nuclear mGlu_5_ signals emanate from pools comprised of both natively expressed nuclear mGlu_5_ and receptors shuttled from the cell surface via endosomes. mGlu_5_ contains nuclear membrane targeting sequences, phosphorylation or glycosylation of which facilitate translocation to nuclear membranes after ligand-mediated receptor internalization ^108^. Endoplasmic reticulum and nuclear mGlu_5_ activates signaling pathways distinct from cell surface receptors, including sustained Ca^2+^ and ERK1/2 signaling ^7,14,21,25,90^. These signaling pathways contribute to pain pathology, as inhibiting glutamate transport into the cell or blocking intracellular mGlu_5_ with cell permeable inhibitors mitigates pain behaviors in animals ^7,14^. While the downstream consequences of endosomal signaling by mGlu_5_ are yet to be fully identified, the contribution of this receptor pool to overall mGlu_5_ signaling is of particular interest, especially given increasing evidence for the role of sustained endosomal GPCR signaling in nociception ^33–37,41,42,46,58^.

Although mGlu_5_ canonically signals through Gα_q_, the receptor is pleiotropically coupled and also recruits Gα_s_ in recombinant cell lines ^109^. Subsequent Gα_s_ signaling through cAMP stimulation is cell type dependent ^75,109–111^. Here we show that G protein coupling to mGlu_5_ is also spatially dependent in HEK293A cells. mGlu5 couples to Gα_q_ at both the cell surface and endosomes, but recruits Gα_s_ only to the cell surface. Similar location selectivity is also evident for other classes of GPCRs such as the free fatty acid receptor 2 (FFAR2), glucagon like peptide receptor 1 (GLP1-R) and Gastrin-Releasing Peptide Receptor (GRPR), as well as the calcium sensing receptor (CaSR), a closely related Class C GPCR ^82,112–115^. While the mechanisms through which distinct, location-specific G protein coupling occurs remain unknown, it is likely that GPCRs can adopt different conformations with distinct G protein coupling profiles. Indeed, cryo-electron microscopy structures of CaSR have revealed distinct conformations for binding to G_αq_ vs G_αi_ ^116^. The CaSR also traffics to early, late and recycling endosomes, where G_αq_, but not G_αs_ or G_αi_, mediates sustained signaling from endosomal compartments. This observation lends further evidence for the structural basis of location-specific G protein coupling ^112,113^. It remains to be seen if other mGlu_5_ ligands, such as positive allosteric modulator agonists (PAM-agonists) or structurally distinct orthosteric agonists, which stabilize different mGlu_5_ conformations, drive the same G protein selectivity as glutamate ^117,118^. Additionally, the contribution of differential G protein coupling from distinct subcellular locations adds another layer of complexity to biased signaling by mGlu_5_, a phenomenon through which different compounds acting on the same receptor can activate divergent signaling pathways ^119^. While endosomal G_αs_ -coupled receptor signaling has been widely reported, evidence for endosomal G_αq_-coupled receptors is less evident. Questions remain about how G_αq_ signaling is mediated from endosomes, with the absence of the canonical phospholipase C substrates in endosomal membranes ^120^. Studies of G protein recruitment to endosomes are also confounded by recent observations that GPCR activation stimulates intracellular redistribution of G proteins in a manner independent of receptor internalization ^61,121,122^. Additionally, constitutive endocytosis is responsible for the majority of G protein abundance on endosomal compartments ^61,123^. It therefore follows that not all G proteins recruited to endosomes will engage in sustained signaling. Our BRET assays confirm these observations: simple BRET between G protein and endosomal markers revealed a substantial movement of G_αq_ to endosomes upon mGlu_5_ activation. However, in nbBRET we observed only a small amount of G_αq_ recruited to endosomally located mGlu_5_ relative to cell surface mGlu_5._ Under the same conditions, activation of NK_1_R resulted in robust and sustained G protein recruitment, indicating assay sensitivity is sufficient to detect endosomal recruitment. Whether the small endosomal recruitment signal evident for mGlu_5_ is reflective of a small pool of signaling competent mGlu_5_ on endosomes, or is simply an artefact of the relative size of the endosomal vs cell surface receptor pools, remains to be seen.

mGlu_5_ is expressed at all points along the pain neuraxis, both centrally and peripherally, and changes in brain and spinal cord mGlu_5_ expression and its signaling partners have been observed in multiple animal models of pain ^7,10,14,124–130^. Importantly, this includes upregulation of mGlu_5_ localized to nuclear membranes within the spinal cord ^7,14^. Additionally, increases in glutamate concentration following pain initiation, and intrathecal or peripheral administration of mGlu_5_ agonists into rodents, are associated with enhanced sensitivity and hyperalgesia ^87,131–134^. Given the clear role of mGlu_5_ activation in pain etiology, attention has turned to inhibitors as potential therapeutics ^135^. mGlu_5_ NAMs from a range of chemical scaffolds are efficacious analgesics in a broad range of acute, inflammatory and neuropathic pain models *in vivo* ^135,136^. Inhibiting intracellular mGlu_5_ signaling plays a role in analgesic efficacy, as membrane impermeable mGlu_5_ antagonists have little effect on pain behaviors when compared to NAMs ^7,14^. mGlu_5_ NAMs have logP values consistent with membrane permeable compounds, although it remains unclear if the accumulation of NAMs in cells reaches therapeutically relevant doses. Additionally, inhibition of glutamate transport into the cell via blockade of glutamate transporters like EAAT3 mimics the analgesic efficacy of NAM-induced mGlu_5_ inhibition ^7,14,137^. mGlu_5_ NAMs also inhibit mGlu_5_ internalization in recombinant cell systems ^96^. If such a mechanism was also present *in vivo*, mGlu_5_ NAMs may promote analgesia by preventing mGlu_5_ internalization, endosomal G protein coupling and subsequent nuclear signaling events associated with pain. Although promising in animal models of pain, mGlu_5_ NAMs have failed to translate clinically ^135^. The mGlu_5_ NAMs raseglurant, AZD2066 and fenobam have all undergone clinical trials for pain, but not progressed beyond Phase II ^135^. Adverse side effects such as cognitive impairment, psychotomimetic effects and tolerance have been observed in preclinical models, and clinical trials have revealed adverse effects such as liver toxicity, dizziness, nausea and anxiety ^135^. On top of safety concerns, poor efficacy and high inter-individual variability have limited the clinical utility of mGlu_5_ NAMs ^53^. While the therapeutic utility of mGlu_5_ NAMs for treating pain is clear, it is also evident that adaptations to current paradigms are needed to mitigate the efficacy and safety concerns that currently hamper clinical translation. In our view, favoring endosomal delivery of drugs provides a more efficacious and sustained approach to pain treatment using mGlu_5_ NAMs.

Drug delivery directly to intracellular GPCRs is gaining traction as a novel approach for the treatment of pain. Along with mGlu_5_, a number of other pain-associated GPCRs signal from intracellular compartments, including neurokinin 1 receptor (NK1R), protease activated receptor 2 (PAR2), calcitonin-like receptor (CLR), GRPR, and the delta and mu opioid receptors (DOR and MOR) ^33–36,41,42,46,58,82,138^. Several approaches have been utilized to target endosomal signaling of these GPCRs. Cholestanol conjugates of NK1R, CLR and PAR2 antagonists accumulate within endosomal membranes, inhibit endosomal GPCR signaling and are longer lasting and more efficacious analgesics than conventional antagonists ^33–36,46^. Biomaterial nanostars containing the NK1R antagonist aprepitant accumulate in spinal cord neuron endosomes, inhibit NK1R endosomal signaling and are superior to free aprepitant with regards to analgesic efficacy ^45^. Similarly, nanoparticle encapsulation of NK1R or CLR antagonists, or DOR agonists, results in drug delivery directly to endosomes, where the acidic environment stimulates drug release, resulting in modulation of sustained GPCR signaling and prolonged analgesia ^36,37,42^. In the current study, we encapsulated the mGlu_5_ NAM VU0366058 in DIPMA nanoparticles to enhance intracellular delivery ^37,54,139^. DIPMA nanoparticles associated with early endosomal markers in HEK293A cells and showed punctate localization consistent with uptake into endosomal compartments in spinal cord neurons. Encapsulation in DIPMA nanoparticles enhanced the inhibitory properties of VU0366058 on sustained cytoplasmic and nuclear mGlu_5_ signaling, reduced the excitability of nociceptive spinal cord neurons and provided superior and longer lasting analgesia in models of acute, neuropathic and inflammatory pain when compared to both free VU0366058 and the clinical candidate mGlu_5_ NAM fenobam. Encapsulation of mGlu_5_ NAMs may offer two advantages in the treatment of pain; first, delivery to endosomes and other intracellular compartments will selectively inhibit sustained intracellular signaling pathways associated with pain pathology. Secondly, by delivering directly into the cell, the use of lower doses may be feasible, potentially bypassing safety concerns associated with mGlu_5_ NAMs. Although enhancing blockade of intracellular mGlu_5_ provides better analgesic efficacy in animal models, it remains to be seen if cell surface vs intracellular mGlu_5_ pools differentially mediate adverse effects ^7,14^. While more studies are needed with different mGlu_5_ NAM chemotypes, our work highlights the feasibility of endosomal mGlu_5_ NAM delivery for effective pain relief.

In conclusion, the current study demonstrates for the first time that mGlu_5_, a Class C GPCR, continues to signal from endosomes upon internalization, with a distinct G protein selectivity when compared to cell surface signaling. Targeting endosomal signaling through nanoparticle encapsulation, VU0366058 reduced sustained signaling across pain-relevant pathways, suppressed neuronal excitability in nociceptive spinal cord circuitry and enhanced the analgesic properties when compared to free drug and fenobam. While the current study focused on endosomal and intracellular mGlu_5_ signaling in the context of pain, it remains to be seen if such pathways are relevant in other pathological conditions in which mGlu_5_ plays a role, such as Alzheimer’s disease, Parkinson’s disease, Fragile X syndrome and multiple neuropsychiatric conditions. By understanding localized signaling in both physiological and pathophysiological conditions, we may successfully develop targeted therapies to fine tune mGlu_5_ activity in a location-dependent manner, overcoming barriers to clinical translation that currently hamper mGlu_5_ drug discovery.

## Acknowledgements

The authors thank Professor Nevin A. Lambert (Augusta University, USA) for providing mini-G-protein-Venus/NLuc (mG_s_, mG_q/11,_ mG_i/0_ or mG_12/13_) and CAAX/FYVE-LgBit DNA and Professor Kevin Pfleger (University of Western Australia, Australia) for providing mGlu_5_-Rluc8 constructs

## Author contributions

Participated in research design: JSR, PRG, DPP, NAV, MW, SH, KJG, WI.

Conducted experiments: JRS, PRG, RL, JK, SH, RP, YZ, WI, KO.

Performed data analysis: JSR, PRG, JK, SH, YZ, RP, WI.

Contributed to writing or critical assessment of the paper: JSR, SH, PRG, DPP, NAV, KJG, WI

## References

1. Gregory KJ., Goudet C. International Union of Basic and Clinical Pharmacology. CXI. Pharmacology, Signaling, and Physiology of Metabotropic Glutamate Receptors. Pharmacol Rev 2021;73(1):521–69. Doi: 10.1124/pr.119.019133.

2. Traynelis SF., Wollmuth LP., McBain CJ., Menniti FS., Vance KM., Ogden KK., Hansen KB., Yuan H., Myers SJ., Dingledine R. Glutamate receptor ion channels: structure, regulation, and function. Pharmacol Rev 2010;62(3):405–96. Doi: 10.1124/pr.109.002451.

3. Sugiyama H., Ito I., Hirono C. A new type of glutamate receptor linked to inositol phospholipid metabolism. Nature 1987;325(6104):531–3. Doi: 10.1038/325531a0.

4. Vidnyánszky Z., Hámori J., Négyessy L., Rüegg D., Knöpfel T., Kuhn R., Görcs TJ. Cellular and subcellular localization of the mGluR5a metabotropic glutamate receptor in rat spinal cord. Neuroreport 1994;6(1):209–13. Doi: 10.1097/00001756-199412300-00053.

5. Alvarez FJ., Villalba RM., Carr PA., Grandes P., Somohano PM. Differential distribution of metabotropic glutamate receptors 1a, 1b, and 5 in the rat spinal cord. J Comp Neurol 2000;422(3):464–87. Doi: 10.1002/1096-9861(20000703)422:3%3C464::aid-cne11%3E3.0.co;2-%23.

6. Neugebauer V. Metabotropic glutamate receptors--important modulators of nociception and pain behavior. Pain 2002;98(1–2):1–8. Doi: 10.1016/s0304-3959(02)00140-9.

7. Vincent K., Cornea VM., Jong Y-JI., Laferrière A., Kumar N., Mickeviciute A., Fung JST., Bandegi P., Ribeiro-da-Silva A., O’Malley KL., Coderre TJ. Intracellular mGluR5 plays a critical role in neuropathic pain. Nat Commun 2016;7(1):10604. Doi: 10.1038/ncomms10604.

8. Honda K., Shinoda M., Kondo M., Shimizu K., Yonemoto H., Otsuki K., Akasaka R., Furukawa A., Iwata K. Sensitization of TRPV1 and TRPA1 via peripheral mGluR5 signaling contributes to thermal and mechanical hypersensitivity. Pain 2017;158(9):1754–64. Doi: 10.1097/j.pain.0000000000000973.

9. Lax NC., George DC., Ignatz C., Kolber BJ. The mGluR5 Antagonist Fenobam Induces Analgesic Conditioned Place Preference in Mice with Spared Nerve Injury. PLOS ONE 2014;9(7):e103524. Doi: 10.1371/journal.pone.0103524.

10. Niu Y., Zeng X., Zhao L., Zhou Y., Qin G., Zhang D., Fu Q., Zhou J., Chen L. Metabotropic glutamate receptor 5 regulates synaptic plasticity in a chronic migraine rat model through the PKC/NR2B signal. J Headache Pain 2020;21(1):139. Doi: 10.1186/s10194-020-01206-2.

11. Xiao XL., Ma DL., Wu J., Tang F-R. Metabotropic glutamate receptor 5 (mGluR5) regulates proliferation and differentiation of neuronal progenitors in the developmental hippocampus. Brain Res 2013;1493:1–12. Doi: 10.1016/j.brainres.2012.11.015.

12. Eng AG., Kelver DA., Hedrick TP., Swanson GT. Transduction of group I mGluR-mediated synaptic plasticity by β-arrestin2 signalling. Nat Commun 2016;7(1):13571. Doi: 10.1038/ncomms13571.

13. Xie J-D., Chen S-R., Pan H-L. Presynaptic mGluR5 receptor controls glutamatergic input through protein kinase C–NMDA receptors in paclitaxel-induced neuropathic pain. J Biol Chem 2017;292(50):20644–54. Doi: 10.1074/jbc.M117.818476.

14. Vincent K., Wang SF., Laferrière A., Kumar N., Coderre TJ. Spinal intracellular metabotropic glutamate receptor 5 (mGluR5) contributes to pain and c-fos expression in a rat model of inflammatory pain. Pain 2017;158(4):705–16. Doi: 10.1097/j.pain.0000000000000823.

15. Peterson CD., Kitto KF., Akgün E., Lunzer MM., Riedl MS., Vulchanova L., Wilcox GL., Portoghese PS., Fairbanks CA. A bivalent ligand that activates mu opioid receptor and antagonizes mGluR5 receptor reduces neuropathic pain in mice. Pain 2017;158(12):2431–41. Doi: 10.1097/j.pain.0000000000001050.

16. Petrenko AB., Shimoji K. A possible role for glutamate receptor-mediated excitotoxicity in chronic pain. J Anesth 2001;15(1):39–48. Doi: 10.1007/s005400170050.

17. Zhu CZ., Wilson SG., Mikusa JP., Wismer CT., Gauvin DM., Lynch JJ., Wade CL., Decker MW., Honore P. Assessing the role of metabotropic glutamate receptor 5 in multiple nociceptive modalities. Eur J Pharmacol 2004;506(2):107–18. Doi: 10.1016/j.ejphar.2004.11.005.

18. Smeester BA., Lunzer MM., Akgün E., Beitz AJ., Portoghese PS. Targeting putative mu opioid/metabotropic glutamate receptor-5 heteromers produces potent antinociception in a chronic murine bone cancer model. Eur J Pharmacol 2014;743:48–52. Doi: 10.1016/j.ejphar.2014.09.008.

19. Akgün E., Javed MI., Lunzer MM., Smeester BA., Beitz AJ., Portoghese PS. Ligands that interact with putative MOR-mGluR5 heteromer in mice with inflammatory pain produce potent antinociception. Proc Natl Acad Sci U S A 2013;110(28):11595–9. Doi: 10.1073/pnas.1305461110.

20. O’Malley KL., Jong Y-JI., Gonchar Y., Burkhalter A., Romano C. Activation of metabotropic glutamate receptor mGlu5 on nuclear membranes mediates intranuclear Ca2+ changes in heterologous cell types and neurons. J Biol Chem 2003;278(30):28210–9. Doi: 10.1074/jbc.M300792200.

21. Jong Y-JI., Kumar V., Kingston AE., Romano C., O’Malley KL. Functional Metabotropic Glutamate Receptors on Nuclei from Brain and Primary Cultured Striatal Neurons: ROLE OF TRANSPORTERS IN DELIVERING LIGAND *. J Biol Chem 2005;280(34):30469–80. Doi: 10.1074/jbc.M501775200.

22. Jong Y-JI., Kumar V., O’Malley KL. Intracellular Metabotropic Glutamate Receptor 5 (mGluR5) Activates Signaling Cascades Distinct from Cell Surface Counterparts. J Biol Chem 2009;284(51):35827–38. Doi: 10.1074/jbc.M109.046276.

23. Jong Y-JI., Izumi Y., Harmon SK., Zorumski CF., O’Malley KL. Striatal mGlu5-mediated synaptic plasticity is independently regulated by location-specific receptor pools and divergent signaling pathways. J Biol Chem 2023;0(0). Doi: 10.1016/j.jbc.2023.104949.

24. Kumar V., Fahey PG., Jong Y-JI., Ramanan N., O’Malley KL. Activation of Intracellular Metabotropic Glutamate Receptor 5 in Striatal Neurons Leads to Up-regulation of Genes Associated with Sustained Synaptic Transmission Including Arc/Arg3.1 Protein. J Biol Chem 2012;287(8):5412–25. Doi: 10.1074/jbc.M111.301366.

25. Kumar V., Jong Y-JI., O’Malley KL. Activated nuclear metabotropic glutamate receptor mGlu5 couples to nuclear Gq/11 proteins to generate inositol 1,4,5-trisphosphate-mediated nuclear Ca2+ release. J Biol Chem 2008;283(20):14072–83. Doi: 10.1074/jbc.M708551200.

26. Fourgeaud L., Bessis A-S., Rossignol F., Pin J-P., Olivo-Marin J-C., Hémar A. The metabotropic glutamate receptor mGluR5 is endocytosed by a clathrin-independent pathway. J Biol Chem 2003;278(14):12222–30. Doi: 10.1074/jbc.M205663200.

27. Trivedi RR., Bhattacharyya S. Constitutive internalization and recycling of metabotropic glutamate receptor 5 (mGluR5). Biochem Biophys Res Commun 2012;427(1):185–90. Doi: 10.1016/j.bbrc.2012.09.040.

28. Francesconi A., Kumari R., Zukin RS. Regulation of group I metabotropic glutamate receptor trafficking and signaling by the caveolar/lipid raft pathway. J Neurosci Off J Soc Neurosci 2009;29(11):3590–602. Doi: 10.1523/JNEUROSCI.5824-08.2009.

29. Abreu N., Acosta-Ruiz A., Xiang G., Levitz J. Mechanisms of differential desensitization of metabotropic glutamate receptors. Cell Rep 2021;35(4):109050. Doi: 10.1016/j.celrep.2021.109050.

30. Lee JH., Lee J., Choi KY., Hepp R., Lee J-Y., Lim MK., Chatani-Hinze M., Roche PA., Kim DG., Ahn YS., Kim CH., Roche KW. Calmodulin dynamically regulates the trafficking of the metabotropic glutamate receptor mGluR5. Proc Natl Acad Sci 2008;105(34):12575–80. Doi: 10.1073/pnas.0712033105.

31. Scheefhals N., Catsburg LAE., Westerveld ML., Blanpied TA., Hoogenraad CC., MacGillavry HD. Shank Proteins Couple the Endocytic Zone to the Postsynaptic Density to Control Trafficking and Signaling of Metabotropic Glutamate Receptor 5. Cell Rep 2019;29(2):258–269.e8. Doi: 10.1016/j.celrep.2019.08.102.

32. Mahato PK., Pandey S., Bhattacharyya S. Differential effects of protein phosphatases in the recycling of metabotropic glutamate receptor 5. Neuroscience 2015;306:138–50. Doi: 10.1016/j.neuroscience.2015.08.031.

33. Jensen DD., Lieu T., Halls ML., Veldhuis NA., Imlach WL., Mai QN., Poole DP., Quach T., Aurelio L., Conner J., Herenbrink CK., Barlow N., Simpson JS., Scanlon MJ., Graham B., McCluskey A., Robinson PJ., Escriou V., Nassini R., Materazzi S., Geppetti P., Hicks GA., Christie MJ., Porter CJH., Canals M., Bunnett NW. Neurokinin 1 receptor signaling in endosomes mediates sustained nociception and is a viable therapeutic target for prolonged pain relief. Sci Transl Med 2017;9(392):eaal3447. Doi: 10.1126/scitranslmed.aal3447.

34. Yarwood RE., Imlach WL., Lieu T., Veldhuis NA., Jensen DD., Klein Herenbrink C., Aurelio L., Cai Z., Christie MJ., Poole DP., Porter CJH., McLean P., Hicks GA., Geppetti P., Halls ML., Canals M., Bunnett NW. Endosomal signaling of the receptor for calcitonin gene-related peptide mediates pain transmission. Proc Natl Acad Sci 2017;114(46):12309–14. Doi: 10.1073/pnas.1706656114.

35. Jimenez-Vargas NN., Pattison LA., Zhao P., Lieu T., Latorre R., Jensen DD., Castro J., Aurelio L., Le GT., Flynn B., Herenbrink CK., Yeatman HR., Edgington-Mitchell L., Porter CJH., Halls ML., Canals M., Veldhuis NA., Poole DP., McLean P., Hicks GA., Scheff N., Chen E., Bhattacharya A., Schmidt BL., Brierley SM., Vanner SJ., Bunnett NW. Protease-activated receptor-2 in endosomes signals persistent pain of irritable bowel syndrome. Proc Natl Acad Sci 2018;115(31):E7438–47. Doi: 10.1073/pnas.1721891115.

36. Jimenez-Vargas NN., Gong J., Wisdom MJ., Jensen DD., Latorre R., Hegron A., Teng S., DiCello JJ., Rajasekhar P., Veldhuis NA., Carbone SE., Yu Y., Lopez-Lopez C., Jaramillo-Polanco J., Canals M., Reed DE., Lomax AE., Schmidt BL., Leong KW., Vanner SJ., Halls ML., Bunnett NW., Poole DP. Endosomal signaling of delta opioid receptors is an endogenous mechanism and therapeutic target for relief from inflammatory pain. Proc Natl Acad Sci 2020;117(26):15281–92. Doi: 10.1073/pnas.2000500117.

37. Ramírez-García PD., Retamal JS., Shenoy P., Imlach W., Sykes M., Truong N., Constandil L., Pelissier T., Nowell CJ., Khor SY., Layani LM., Lumb C., Poole DP., Lieu T., Stewart GD., Mai QN., Jensen DD., Latorre R., Scheff NN., Schmidt BL., Quinn JF., Whittaker MR., Veldhuis NA., Davis TP., Bunnett NW. A pH-responsive nanoparticle targets the neurokinin 1 receptor in endosomes to prevent chronic pain. Nat Nanotechnol 2019;14(12):1150–9. Doi: 10.1038/s41565-019-0568-x.

38. Blythe EE., von Zastrow M. β-Arrestin-independent endosomal cAMP signaling by a polypeptide hormone GPCR. Nat Chem Biol 2024;20(3):323–32. Doi: 10.1038/s41589-023-01412-4.

39. Ismail S., Gherardi M-J., Froese A., Zanoun M., Gigoux V., Clerc P., Gaits-Iacovoni F., Steyaert J., Nikolaev VO., Fourmy D. Internalized Receptor for Glucose-dependent Insulinotropic Peptide stimulates adenylyl cyclase on early endosomes. Biochem Pharmacol 2016;120:33–45. Doi: 10.1016/j.bcp.2016.09.009.

40. Peach CJ., Tonello R., Damo E., Gomez K., Calderon-Rivera A., Bruni R., Bansia H., Maile L., Manu A-M., Hahn H., Thomsen ARB., Schmidt BL., Davidson S., Georges A des., Khanna R., Bunnett NW. NEUROPILIN-1 INHIBITION SUPPRESSES NERVE-GROWTH FACTOR SIGNALING AND NOCICEPTION IN PAIN MODELS. J Clin Invest 2024. Doi: 10.1172/JCI183873.

41. Hegron A., Peach CJ., Tonello R., Seemann P., Teng S., Latorre R., Huebner H., Weikert D., Rientjes J., Veldhuis NA., Poole DP., Jensen DD., Thomsen ARB., Schmidt BL., Imlach WL., Gmeiner P., Bunnett NW. Therapeutic antagonism of the neurokinin 1 receptor in endosomes provides sustained pain relief. Proc Natl Acad Sci 2023;120(22):e2220979120. Doi: 10.1073/pnas.2220979120.

42. De Logu F., Nassini R., Hegron A., Landini L., Jensen DD., Latorre R., Ding J., Marini M., Souza Monteiro de Araujo D., Ramírez-Garcia P., Whittaker M., Retamal J., Titiz M., Innocenti A., Davis TP., Veldhuis N., Schmidt BL., Bunnett NW., Geppetti P. Schwann cell endosome CGRP signals elicit periorbital mechanical allodynia in mice. Nat Commun 2022;13(1):646. Doi: 10.1038/s41467-022-28204-z.

43. Kwon Y., Mehta S., Clark M., Walters G., Zhong Y., Lee HN., Sunahara RK., Zhang J. Non-canonical β-adrenergic activation of ERK at endosomes. Nature 2022;611(7934):173–9. Doi: 10.1038/s41586-022-05343-3.

44. Irannejad R., Tomshine JC., Tomshine JR., Chevalier M., Mahoney JP., Steyaert J., Rasmussen SGF., Sunahara RK., El-Samad H., Huang B., Von Zastrow M. Conformational biosensors reveal GPCR signalling from endosomes. Nature 2013;495(7442):534–8. Doi: 10.1038/nature12000.

45. Latorre R., Ramírez-Garcia PD., Hegron A., Grace JL., Retamal JS., Shenoy P., Tran M., Aurelio L., Flynn B., Poole DP., Klein-Cloud R., Jensen DD., Davis TP., Schmidt BL., Quinn JF., Whittaker MR., Veldhuis NA., Bunnett NW. Sustained endosomal release of a neurokinin-1 receptor antagonist from nanostars provides long-lasting relief of chronic pain. Biomaterials 2022;285:121536. Doi: 10.1016/j.biomaterials.2022.121536.

46. Mai QN., Shenoy P., Quach T., Retamal JS., Gondin AB., Yeatman HR., Aurelio L., Conner JW., Poole DP., Canals M., Nowell CJ., Graham B., Davis TP., Briddon SJ., Hill SJ., Porter CJH., Bunnett NW., Halls ML., Veldhuis NA. A lipid-anchored neurokinin 1 receptor antagonist prolongs pain relief by a three-pronged mechanism of action targeting the receptor at the plasma membrane and in endosomes. J Biol Chem 2021;296. Doi: 10.1016/j.jbc.2021.100345.

47. Jong Y-JI., Harmon SK., O’Malley KL. Location and Cell-Type-Specific Bias of Metabotropic Glutamate Receptor, mGlu5, Negative Allosteric Modulators. ACS Chem Neurosci 2019. Doi: 10.1021/acschemneuro.9b00415.

48. Montana MC., Conrardy BA., Cavallone LF., Kolber BJ., Rao LK., Greco SC., Gereau RW. Metabotropic glutamate receptor 5 antagonism with fenobam: examination of analgesic tolerance and side effect profile in mice. Anesthesiology 2011;115(6):1239–50. Doi: 10.1097/ALN.0b013e318238c051.

49. Sevostianova N., Danysz W. Analgesic effects of mGlu1 and mGlu5 receptor antagonists in the rat formalin test. Neuropharmacology 2006;51(3):623–30. Doi: 10.1016/j.neuropharm.2006.05.004.

50. Crock LW., Stemler KM., Song DG., Abbosh P., Vogt SK., Qiu C-S., Lai HH., Mysorekar IU., Gereau IV RW. Metabotropic glutamate receptor 5 (mGluR5) regulates bladder nociception. Mol Pain 2012;8:20. Doi: 10.1186/1744-8069-8-20.

51. Notartomaso S., Antenucci N., Mazzitelli M., Rovira X., Boccella S., Ricciardi F., Liberatore F., Gomez-Santacana X., Imbriglio T., Cannella M., Zussy C., Luongo L., Maione S., Goudet C., Battaglia G., Llebaria A., Nicoletti F., Neugebauer V. A ‘double-edged’ role for type-5 metabotropic glutamate receptors in pain disclosed by light-sensitive drugs. eLife 2024;13:e94931. Doi: 10.7554/eLife.94931.

52. Abou Farha K., Bruggeman R., Baljé-Volkers C. Metabotropic glutamate receptor 5 negative modulation in phase I clinical trial: potential impact of circadian rhythm on the neuropsychiatric adverse reactions-do hallucinations matter? ISRN Psychiatry 2014;2014:652750. Doi: 10.1155/2014/652750.

53. Cavallone LF., Montana MC., Frey K., Kallogjeri D., Wages JM., Rodebaugh TL., Doshi T., Kharasch ED., Gereau RW. The metabotropic glutamate receptor 5 negative allosteric modulator fenobam: pharmacokinetics, side effects, and analgesic effects in healthy human subjects. Pain 2020;161(1):135–46. Doi: 10.1097/j.pain.0000000000001695.

54. Mueller R., Dawson ES., Meiler J., Rodriguez AL., Chauder BA., Bates BS., Felts AS., Lamb JP., Menon UN., Jadhav SB., Kane AS., Jones CK., Gregory KJ., Niswender CM., Conn PJ., Olsen CM., Winder DG., Emmitte KA., Lindsley CW. Discovery of 2-(2-benzoxazoyl amino)-4-aryl-5-cyanopyrimidine as negative allosteric modulators (NAMs) of metabotropic glutamate receptor 5 (mGlu(5)): from an artificial neural network virtual screen to an in vivo tool compound. ChemMedChem 2012;7(3):406–14. Doi: 10.1002/cmdc.201100510.

55. Noetzel MJ., Rook JM., Vinson PN., Cho HP., Days E., Zhou Y., Rodriguez AL., Lavreysen H., Stauffer SR., Niswender CM., Xiang Z., Daniels JS., Jones CK., Lindsley CW., Weaver CD., Conn PJ. Functional impact of allosteric agonist activity of selective positive allosteric modulators of metabotropic glutamate receptor subtype 5 in regulating central nervous system function. Mol Pharmacol 2012;81(2):120–33. Doi: 10.1124/mol.111.075184.

56. Harvey CD., Ehrhardt AG., Cellurale C., Zhong H., Yasuda R., Davis RJ., Svoboda K. A genetically encoded fluorescent sensor of ERK activity. Proc Natl Acad Sci U S A 2008;105(49):19264–9. Doi: 10.1073/pnas.0804598105.

57. Zhao Y., Araki S., Wu J., Teramoto T., Chang Y-F., Nakano M., Abdelfattah AS., Fujiwara M., Ishihara T., Nagai T., Campbell RE. An expanded palette of genetically encoded Ca^2+^ indicators. Science 2011;333(6051):1888–91. Doi: 10.1126/science.1208592.

58. Latorre R., Hegron A., Peach CJ., Teng S., Tonello R., Retamal JS., Klein-Cloud R., Bok D., Jensen DD., Gottesman-Katz L., Rientjes J., Veldhuis NA., Poole DP., Schmidt BL., Pothoulakis CH., Rankin C., Xie Y., Koon HW., Bunnett NW. Mice expressing fluorescent PAR2 reveal that endocytosis mediates colonic inflammation and pain. Proc Natl Acad Sci U S A 2022;119(6):e2112059119. Doi: 10.1073/pnas.2112059119.

59. Wan Q., Okashah N., Inoue A., Nehmé R., Carpenter B., Tate CG., Lambert NA. Mini G protein probes for active G protein-coupled receptors (GPCRs) in live cells. J Biol Chem 2018;293(19):7466–73. Doi: 10.1074/jbc.RA118.001975.

60. Herskovits J., Burgess C., Obar R., Vallee R. Effects of mutant rat dynamin on endocytosis. J Cell Biol 1993;122(3):565–78. Doi: 10.1083/jcb.122.3.565.

61. Martin BR., Lambert NA. Activated G Protein Gαs Samples Multiple Endomembrane Compartments♦. J Biol Chem 2016;291(39):20295–302. Doi: 10.1074/jbc.M116.729731.

62. Schindelin J., Arganda-Carreras I., Frise E., Kaynig V., Longair M., Pietzsch T., Preibisch S., Rueden C., Saalfeld S., Schmid B., Tinevez J-Y., White DJ., Hartenstein V., Eliceiri K., Tomancak P., Cardona A. Fiji: an open-source platform for biological-image analysis. Nat Methods 2012;9(7):676–82. Doi: 10.1038/nmeth.2019.

63. Kilkenny C., Browne WJ., Cuthill IC., Emerson M., Altman DG. Improving bioscience research reporting: The ARRIVE guidelines for reporting animal research. J Pharmacol Pharmacother 2010;1(2):94–9. Doi: 10.4103/0976-500X.72351.

64. Dixon AS., Schwinn MK., Hall MP., Zimmerman K., Otto P., Lubben TH., Butler BL., Binkowski BF., Machleidt T., Kirkland TA., Wood MG., Eggers CT., Encell LP., Wood KV. NanoLuc Complementation Reporter Optimized for Accurate Measurement of Protein Interactions in Cells. ACS Chem Biol 2016;11(2):400–8. Doi: 10.1021/acschembio.5b00753.

65. Retamal JS., Grace MS., Dill LK., Ramirez-Garcia P., Peng S., Gondin AB., Bennetts F., Alvi S., Rajasekhar P., Almazi JG., Carbone SE., Bunnett NW., Davis TP., Veldhuis NA., Poole DP., McIntyre P. Serotonin-induced vascular permeability is mediated by transient receptor potential vanilloid 4 in the airways and upper gastrointestinal tract of mice. Lab Invest 2021;101(7):851–64. Doi: 10.1038/s41374-021-00593-7.

66. Peng S., Grace MS., Gondin AB., Retamal JS., Dill L., Darby W., Bunnett NW., Abogadie FC., Carbone SE., Tigani T., Davis TP., Poole DP., Veldhuis NA., McIntyre P. The transient receptor potential vanilloid 4 (TRPV4) ion channel mediates protease activated receptor 1 (PAR1)-induced vascular hyperpermeability. Lab Investig J Tech Methods Pathol 2020;100(8):1057–67. Doi: 10.1038/s41374-020-0430-7.

67. Gregory KJ., Noetzel MJ., Rook JM., Vinson PN., Stauffer SR., Rodriguez AL., Emmitte KA., Zhou Y., Chun AC., Felts AS., Chauder BA., Lindsley CW., Niswender CM., Conn PJ. Investigating Metabotropic Glutamate Receptor 5 Allosteric Modulator Cooperativity, Affinity, and Agonism: Enriching Structure-Function Studies and Structure-Activity Relationships. Mol Pharmacol 2012;82(5):860–75. Doi: 10.1124/mol.112.080531.

68. Imlach WL., Bhola RF., Mohammadi SA., Christie MJ. Glycinergic dysfunction in a subpopulation of dorsal horn interneurons in a rat model of neuropathic pain. Sci Rep 2016;6:37104. Doi: 10.1038/srep37104.

69. Manders EMM., Verbeek FJ., Aten JA. Measurement of co-localization of objects in dual-colour confocal images. J Microsc 1993;169(3):375–82. Doi: 10.1111/j.1365-2818.1993.tb03313.x.

70. Cichon J., Sun L., Yang G. Spared Nerve Injury Model of Neuropathic Pain in Mice. Bio-Protoc 2018;8(6):e2777. Doi: 10.21769/bioprotoc.2777.

71. Bhattacharya M., Babwah AV., Godin C., Anborgh PH., Dale LB., Poulter MO., Ferguson SSG. Ral and Phospholipase D2-Dependent Pathway for Constitutive Metabotropic Glutamate Receptor Endocytosis. J Neurosci 2004;24(40):8752–61. Doi: 10.1523/JNEUROSCI.3155-04.2004.

72. Gales C., Rebois RV., Hogue M., Trieu P., Breit A., Hebert TE., Bouvier M. Real-time monitoring of receptor and G-protein interactions in living cells. Nat Methods 2005;2(3):177–84. Doi: 10.1038/nmeth743.

73. Harper CB., Martin S., Nguyen TH., Daniels SJ., Lavidis NA., Popoff MR., Hadzic G., Mariana A., Chau N., McCluskey A., Robinson PJ., Meunier FA. Dynamin inhibition blocks botulinum neurotoxin type A endocytosis in neurons and delays botulism. J Biol Chem 2011;286(41):35966–76. Doi: 10.1074/jbc.M111.283879.

74. Joly C., Gomeza J., Brabet I., Curry K., Bockaert J., Pin JP. Molecular, functional, and pharmacological characterization of the metabotropic glutamate receptor type 5 splice variants: comparison with mGluR1. J Neurosci 1995;15(5):3970–81. Doi: 10.1523/JNEUROSCI.15-05-03970.1995.

75. Francesconi A., Duvoisin RM. Role of the second and third intracellular loops of metabotropic glutamate receptors in mediating dual signal transduction activation. J Biol Chem 1998;273(10):5615–24. Doi: 10.1074/jbc.273.10.5615.

76. Gegelashvili G., Bjerrum OJ. Glutamate Transport System as a Novel Therapeutic Target in Chronic Pain: Molecular Mechanisms and Pharmacology. Adv Neurobiol 2017;16:225–53. Doi: 10.1007/978-3-319-55769-4_11.

77. Purgert CA., Izumi Y., Jong Y-JI., Kumar V., Zorumski CF., O’Malley KL. Intracellular mGluR5 Can Mediate Synaptic Plasticity in the Hippocampus. J Neurosci 2014;34(13):4589–98. Doi: 10.1523/JNEUROSCI.3451-13.2014.

78. van der Bliek A., Redelmeier T., Damke H., Tisdale E., Meyerowitz E., Schmid S. Mutations in human dynamin block an intermediate stage in coated vesicle formation. J Cell Biol 1993;122(3):553–63. Doi: 10.1083/jcb.122.3.553.

79. Toki H., Namikawa K., Su Q., Kiryu-Seo S., Sato K., Kiyama H. Enhancement of Extracellular Glutamate Scavenge System in Injured Motoneurons. J Neurochem 1998;71(3):913–9. Doi: 10.1046/j.1471-4159.1998.71030913.x.

80. Kelner A., Leitão N., Chabaud M., Charpentier M., de Carvalho-Niebel F. Dual Color Sensors for Simultaneous Analysis of Calcium Signal Dynamics in the Nuclear and Cytoplasmic Compartments of Plant Cells. Front Plant Sci 2018;9:245. Doi: 10.3389/fpls.2018.00245.

81. Seidler NW., Jona I., Vegh M., Martonosi A. Cyclopiazonic acid is a specific inhibitor of the Ca2+-ATPase of sarcoplasmic reticulum. J Biol Chem 1989;264(30):17816–23.

82. Santibañez JR., Bok D., Teng S., Bhansali D., Ferreira M de A., Tonello R., Peach CJ., Latorre R., Thanigai GS., Leong KW., Jensen DD. Characterization and targeting of the endosomal signaling of the gastrin releasing peptide receptor in pruritus 2025:2025.03.17.643743. Doi: 10.1101/2025.03.17.643743.

83. Lax NC., George DC., Ignatz C., Kolber BJ. The mGluR5 antagonist fenobam induces analgesic conditioned place preference in mice with spared nerve injury. PloS One 2014;9(7):e103524. Doi: 10.1371/journal.pone.0103524.

84. Jacob W., Gravius A., Pietraszek M., Nagel J., Belozertseva I., Shekunova E., Malyshkin A., Greco S., Barberi C., Danysz W. The anxiolytic and analgesic properties of fenobam, a potent mGlu5 receptor antagonist, in relation to the impairment of learning. Neuropharmacology 2009;57(2):97–108. Doi: 10.1016/j.neuropharm.2009.04.011.

85. Jin Y-H., Takemura M., Furuyama A., Yonehara N. Peripheral Glutamate Receptors Are Required for Hyperalgesia Induced by Capsaicin. Pain Res Treat 2012;2012:915706. Doi: 10.1155/2012/915706.

86. Medvedeva YV., Kim M-S., Usachev YM. Mechanisms of Prolonged Presynaptic Ca2+ Signaling and Glutamate Release Induced by TRPV1 Activation in Rat Sensory Neurons. J Neurosci 2008;28(20):5295–311. Doi: 10.1523/JNEUROSCI.4810-07.2008.

87. Kumar N., Laferriere A., Yu JSC., Poon T., Coderre TJ. Metabotropic glutamate receptors (mGluRs) regulate noxious stimulus-induced glutamate release in spinal cord dorsal horn of rats with neuropathic and inflammatory pain. J Neurochem 2010;114(1):281–90. Doi: 10.1111/j.1471-4159.2010.06761.x.

88. Soliman AC., Yu JSC., Coderre TJ. mGlu and NMDA receptor contributions to capsaicin-induced thermal and mechanical hypersensitivity. Neuropharmacology 2005;48(3):325–32. Doi: 10.1016/j.neuropharm.2004.10.014.

89. Decosterd I., Woolf CJ. Spared nerve injury: an animal model of persistent peripheral neuropathic pain. PAIN 2000;87(2):149. Doi: 10.1016/S0304-3959(00)00276-1.

90. Jong Y-JI., Harmon SK., O’Malley KL. Activation of Endoplasmic Reticulum-Localized Metabotropic Glutamate Receptor 5 (mGlu5) Triggers Calcium Release Distinct from Cell Surface Counterparts in Striatal Neurons. Biomolecules 2025;15(4):552. Doi: 10.3390/biom15040552.

91. Jong Y-JI., Sergin I., Purgert CA., O’Malley KL. Location-Dependent Signaling of the Group 1 Metabotropic Glutamate Receptor mGlu5. Mol Pharmacol 2014;86(6):774–85. Doi: 10.1124/mol.114.094763.

92. Manengu C., Zhu C-H., Zhang G-D., Tian M-M., Lan X-B., Tao L-J., Ma L., Liu Y., Yu J-Q., Liu N. Metabotropic Glutamate Receptor 5: A Potential Target for Neuropathic Pain Treatment. Curr Neuropharmacol 2025;23(3):276–94. Doi: 10.2174/1570159X23666241011163035.

93. Ribeiro FM., Ferreira LT., Paquet M., Cregan T., Ding Q., Gros R., Ferguson SS. Phosphorylation-independent regulation of metabotropic glutamate receptor 5 desensitization and internalization by G protein-coupled receptor kinase 2 in neurons. J Biol Chem 2009;284(35):23444–53. Doi: 10.1074/jbc.M109.000778.

94. Cimadevila M., Liu J., Maurel D., Brabet I., Hoscar M., Drube J., Hoffmann C., Inoue A., Rondard P., Lafon PA., Prézeau L., Pin JP. Non-canonical internalization mechanisms of mGlu receptors 2025:2025.02.13.638043. Doi: 10.1101/2025.02.13.638043.

95. Arsova A., Møller TC., Hellyer SD., Vedel L., Foster SR., Hansen JL., Bräuner-Osborne H., Gregory KJ. Positive Allosteric Modulators of Metabotropic Glutamate Receptor 5 as Tool Compounds to Study Signaling Bias. Mol Pharmacol 2021;99(5):328–41. Doi: 10.1124/molpharm.120.000185.

96. Arsova A., Møller TC., Vedel L., Hansen JL., Foster SR., Gregory KJ., Bräuner-Osborne H. Detailed in vitro pharmacological characterization of clinically tested negative allosteric modulators of the metabotropic glutamate receptor 5 (mGlu5). Mol Pharmacol 2020. Doi: 10.1124/mol.119.119032.

97. Ojha P., Pal S., Bhattacharyya S. Regulation of Metabotropic Glutamate Receptor Internalization and Synaptic AMPA Receptor Endocytosis by the Postsynaptic Protein Norbin. J Neurosci 2022;42(5):731–48. Doi: 10.1523/jneurosci.1037-21.2021.

98. Kitano J., Kimura K., Yamazaki Y., Soda T., Shigemoto R., Nakajima Y., Nakanishi S. Tamalin, a PDZ domain-containing protein, links a protein complex formation of group 1 metabotropic glutamate receptors and the guanine nucleotide exchange factor cytohesins. J Neurosci 2002;22(4):1280–9. Doi: 10.1523/JNEUROSCI.22-04-01280.2002.

99. Ko SJ., Isozaki K., Kim I., Lee JH., Cho HJ., Sohn SY., Oh SR., Park S., Kim DG., Kim CH., Roche KW. PKC Phosphorylation Regulates mGluR5 Trafficking by Enhancing Binding of Siah-1A. J Neurosci 2012;32(46):16391–401. Doi: 10.1523/JNEUROSCI.1964-12.2012.

100. Toledano-Zaragoza A., Enriquez-Zarralanga V., Naya-Forcano S., Briz V., Alfaro-Ruíz R., Parra-Martínez M., Mitroi DN., Luján R., Esteban JA., Ledesma MD. Enhanced mGluR5 intracellular activity causes psychiatric alterations in Niemann Pick type C disease. Cell Death Dis 2024;15(10):771. Doi: 10.1038/s41419-024-07158-8.

101. Thomsen ARB., Plouffe B., Cahill TJ., Shukla AK., Tarrasch JT., Dosey AM., Kahsai AW., Strachan RT., Pani B., Mahoney JP., Huang L., Breton B., Heydenreich FM., Sunahara RK., Skiniotis G., Bouvier M., Lefkowitz RJ. GPCR-G Protein-β-Arrestin Super-Complex Mediates Sustained G Protein Signaling. Cell 2016;166(4):907–19. Doi: 10.1016/j.cell.2016.07.004.

102. Cahill TJ., Thomsen ARB., Tarrasch JT., Plouffe B., Nguyen AH., Yang F., Huang L-Y., Kahsai AW., Bassoni DL., Gavino BJ., Lamerdin JE., Triest S., Shukla AK., Berger B., Little J., Antar A., Blanc A., Qu C-X., Chen X., Kawakami K., Inoue A., Aoki J., Steyaert J., Sun J-P., Bouvier M., Skiniotis G., Lefkowitz RJ. Distinct conformations of GPCR–β-arrestin complexes mediate desensitization, signaling, and endocytosis. Proc Natl Acad Sci 2017;114(10):2562–7. Doi: 10.1073/pnas.1701529114.

103. Sutkeviciute I., Vilardaga J-P. Structural insights into emergent signaling modes of G protein-coupled receptors. J Biol Chem 2020;295(33):11626–42. Doi: 10.1074/jbc.REV120.009348.

104. Cimadevila M., Liu J., Maurel D., Brabet I., Hoscar M., Drube J., Hoffmann C., Inoue A., Rondard P., Lafon P-A., Prézeau L., Pin J-P. Non-canonical internalization mechanisms of mGlu receptors. Cell Rep 2025;44(8). Doi: 10.1016/j.celrep.2025.116068.

105. Blythe EE., Zastrow M von. A discrete mode of endosomal GPCR signaling that does not require β-arrestin 2022:2022.09.07.506997. Doi: 10.1101/2022.09.07.506997.

106. Teixeira LB., Blouin M-J., Le Gouill C., Picard L-P., Costa-Neto CM., Bouvier M., Parreiras-e-Silva LT. Sustained Gαs signaling mediated by vasopressin type 2 receptors is ligand dependent but endocytosis and β-arrestin independent. Sci Signal 2025;18(874):eadf6206. Doi: 10.1126/scisignal.adf6206.

107. Daly C., Guseinov AA., Hahn H., Wright A., Tikhonova IG., Thomsen ARB., Plouffe B. β-Arrestin-dependent and -independent endosomal G protein activation by the vasopressin type 2 receptor. eLife n.d.;12:RP87754. Doi: 10.7554/eLife.87754.

108. Sergin I., Jong Y-JI., Harmon SK., Kumar V., O’Malley KL. Sequences within the C Terminus of the Metabotropic Glutamate Receptor 5 (mGluR5) Are Responsible for Inner Nuclear Membrane Localization*. J Biol Chem 2017;292(9):3637–55. Doi: 10.1074/jbc.M116.757724.

109. Nasrallah C., Rottier K., Marcellin R., Compan V., Font J., Llebaria A., Pin J-P., Banères J-L., Lebon G. Direct coupling of detergent purified human mGlu5 receptor to the heterotrimeric G proteins Gq and Gs. Sci Rep 2018;8(1):4407. Doi: 10.1038/s41598-018-22729-4.

110. Joly C., Gomeza J., Brabet I., Curry K., Bockaert J., Pin JP. Molecular, functional, and pharmacological characterization of the metabotropic glutamate receptor type 5 splice variants: comparison with mGluR1. J Neurosci Off J Soc Neurosci 1995;15(5 Pt 2):3970–81. Doi: 10.1523/JNEUROSCI.15-05-03970.1995.

111. Francesconi A., Duvoisin RM. Opposing effects of protein kinase C and protein kinase A on metabotropic glutamate receptor signaling: selective desensitization of the inositol trisphosphate/Ca2+ pathway by phosphorylation of the receptor-G protein-coupling domain. Proc Natl Acad Sci U S A 2000;97(11):6185–90.

112. Wyatt RA., Gallagher MT., Zha L., McCabe CJ., Gorvin CM. A calcium-sensing receptor dileucine motif directs internalization to spatially distinct endosomal signaling pathways. iScience 2025;28(6):112651. Doi: 10.1016/j.isci.2025.112651.

113. Gorvin CM., Rogers A., Hastoy B., Tarasov AI., Frost M., Sposini S., Inoue A., Whyte MP., Rorsman P., Hanyaloglu AC., Breitwieser GE., Thakker RV. AP2σ Mutations Impair Calcium-Sensing Receptor Trafficking and Signaling, and Show an Endosomal Pathway to Spatially Direct G-Protein Selectivity. Cell Rep 2018;22(4):1054–66. Doi: 10.1016/j.celrep.2017.12.089.

114. Wright SC., Motso A., Koutsilieri S., Beusch CM., Sabatier P., Berghella A., Blondel-Tepaz É., Mangenot K., Pittarokoilis I., Sismanoglou D-C., Le Gouill C., Olsen JV., Zubarev RA., Lambert NA., Hauser AS., Bouvier M., Lauschke VM. GLP-1R signaling neighborhoods associate with the susceptibility to adverse drug reactions of incretin mimetics. Nat Commun 2023;14(1):6243. Doi: 10.1038/s41467-023-41893-4.

115. Caengprasath N., Gonzalez-Abuin N., Shchepinova M., Ma Y., Inoue A., Tate EW., Frost G., Hanyaloglu AC. Internalization-Dependent Free Fatty Acid Receptor 2 Signaling Is Essential for Propionate-Induced Anorectic Gut Hormone Release. iScience 2020;23(9):101449. Doi: 10.1016/j.isci.2020.101449.

116. He F., Wu C-G., Gao Y., Rahman SN., Zaoralová M., Papasergi-Scott MM., Gu T-J., Robertson MJ., Seven AB., Li L., Mathiesen JM., Skiniotis G. Allosteric modulation and G-protein selectivity of the Ca2+-sensing receptor. Nature 2024;626(8001):1141–8. Doi: 10.1038/s41586-024-07055-2.

117. Cannone G., Berto L., Malhaire F., Ferguson G., Fouillen A., Balor S., Font-Ingles J., Llebaria A., Goudet C., Kotecha A., K R V., Lebon G. Conformational diversity in class C GPCR positive allosteric modulation. Nat Commun 2025;16(1):619. Doi: 10.1038/s41467-024-55439-9.

118. Koehl A., Hu H., Feng D., Sun B., Zhang Y., Robertson MJ., Chu M., Kobilka TS., Laeremans T., Steyaert J., Tarrasch J., Dutta S., Fonseca R., Weis WI., Mathiesen JM., Skiniotis G., Kobilka BK. Structural insights into the activation of metabotropic glutamate receptors. Nature 2019;566(7742):79–84. Doi: 10.1038/s41586-019-0881-4.

119. Trinh PNH., May LT., Leach K., Gregory KJ. Biased agonism and allosteric modulation of metabotropic glutamate receptor 5. Clin Sci Lond Engl 1979 2018;132(21):2323–38. Doi: 10.1042/CS20180374.

120. Daly C., Plouffe B. Gαq signalling from endosomes: A new conundrum. Br J Pharmacol 2025;182(14):3068–89. Doi: 10.1111/bph.16248.

121. Wysolmerski B., Blythe EE., Zastrow M von. Conformational biosensors delineate endosomal G protein regulation by GPCRs 2025:2025.05.12.653522. Doi: 10.1101/2025.05.12.653522.

122. Wright SC., Lukasheva V., Le Gouill C., Kobayashi H., Breton B., Mailhot-Larouche S., Blondel-Tepaz É., Antunes Vieira N., Costa-Neto C., Héroux M., Lambert NA., Parreiras-e-Silva LT., Bouvier M. BRET-based effector membrane translocation assay monitors GPCR-promoted and endocytosis-mediated Gq activation at early endosomes. Proc Natl Acad Sci 2021;118(20):e2025846118. Doi: 10.1073/pnas.2025846118.

123. Jang W., Senarath K., Feinberg G., Lu S., Lambert NA. Visualization of endogenous G proteins on endosomes and other organelles. eLife 2024;13. Doi: 10.7554/eLife.97033.2.

124. Tianxin Z., Zhongyu H., Weishan Z., Can W., Jiahong L., Shuhan W., Runheng Z., Chang Z., Yuxin M. mGluR5 regulates inflammatory pain and pain aversion of CFA mice by mediating ERK/PI3K signaling pathway. Neurosci Lett 2025:138281. Doi: 10.1016/j.neulet.2025.138281.

125. Tian X., Wang W-T., Zhang M-M., Yang Q-Q., Xu Y-L., Wu J-B., Xie X-X., Wang J-Y., Wang J-Y. Red nucleus mGluR1 and mGluR5 facilitate the development of neuropathic pain through stimulating the expressions of TNF-α and IL-1β. Neurochem Int 2024;178:105786. Doi: 10.1016/j.neuint.2024.105786.

126. Niu Y., Zeng X., Qin G., Zhang D., Zhou J., Chen L. Downregulation of metabotropic glutamate receptor 5 alleviates central sensitization by activating autophagy via inhibiting mTOR pathway in a rat model of chronic migraine. Neurosci Lett 2021;743:135552. Doi: 10.1016/j.neulet.2020.135552.

127. Li J-Q., Chen S-R., Chen H., Cai Y-Q., Pan H-L. Regulation of increased glutamatergic input to spinal dorsal horn neurons by mGluR5 in diabetic neuropathic pain. J Neurochem 2010;112(1):162–72. Doi: 10.1111/j.1471-4159.2009.06437.x.

128. Sotgiu ML., Bellomi P., Biella GEM. The mGluR5 selective antagonist 6-methyl-2-(phenylethynyl)-pyridine reduces the spinal neuron pain-related activity in mononeuropathic rats. Neurosci Lett 2003;342(1–2):85–8. Doi: 10.1016/s0304-3940(03)00259-3.

129. Lai C-Y., Hsieh M-C., Ho Y-C., Wang H-H., Chou D., Wen Y-C., Yang P-S., Cheng J-K., Peng H-Y. Spinal RNF20-Mediated Histone H2B Monoubiquitylation Regulates mGluR5 Transcription for Neuropathic Allodynia. J Neurosci 2018;38(43):9160–74. Doi: 10.1523/JNEUROSCI.1069-18.2018.

130. Kartha S., Ghimire P., Winkelstein BA. Inhibiting spinal secretory phospholipase A2 after painful nerve root injury attenuates established pain and spinal neuronal hyperexcitability by altering spinal glutamatergic signaling. Mol Pain 2021;17:17448069211066221. Doi: 10.1177/17448069211066221.

131. Walker K., Reeve A., Bowes M., Winter J., Wotherspoon G., Davis A., Schmid P., Gasparini F., Kuhn R., Urban L. mGlu5 receptors and nociceptive function II. mGlu5 receptors functionally expressed on peripheral sensory neurones mediate inflammatory hyperalgesia. Neuropharmacology 2001;40(1):10–9. Doi: 10.1016/s0028-3908(00)00114-3.

132. Hama AT. Acute activation of the spinal cord metabotropic glutamate subtype-5 receptor leads to cold hypersensitivity in the rat. Neuropharmacology 2003;44(4):423–30. Doi: 10.1016/s0028-3908(03)00026-1.

133. Karim F., Wang C-C., Gereau RW. Metabotropic Glutamate Receptor Subtypes 1 and 5 Are Activators of Extracellular Signal-Regulated Kinase Signaling Required for Inflammatory Pain in Mice. J Neurosci 2001;21(11):3771–9. Doi: 10.1523/JNEUROSCI.21-11-03771.2001.

134. Li Y., Kang J., Xu Y., Li N., Jiao Y., Wang C., Wang C., Wang G., Yu Y., Yuan J., Zhang L. Artesunate Alleviates Paclitaxel-Induced Neuropathic Pain in Mice by Decreasing Metabotropic Glutamate Receptor 5 Activity and Neuroinflammation in Primary Sensory Neurons. Front Mol Neurosci 2022;15:902572. Doi: 10.3389/fnmol.2022.902572.

135. Kos JA., Langiu M., Hellyer SD., Gregory KJ. Pharmacology, Signaling and Therapeutic Potential of Metabotropic Glutamate Receptor 5 Negative Allosteric Modulators. ACS Pharmacol Transl Sci 2024;7(12):3671–90. Doi: 10.1021/acsptsci.4c00213.

136. Pereira V., Goudet C. Emerging Trends in Pain Modulation by Metabotropic Glutamate Receptors. Front Mol Neurosci 2019;11:464. Doi: 10.3389/fnmol.2018.00464.

137. Li X., Yang H., Qian M., Liu H., Zuo S., Liu J-C., Ge W-H., Zhou L. Intracellular metabotropic glutamate receptor 5 in spinal dorsal horn neurons contributes to pain in a mouse model of vincristine-induced neuropathic pain. Neurosci Lett 2025;852:138193. Doi: 10.1016/j.neulet.2025.138193.

138. Stoeber M., Jullié D., Lobingier BT., Laeremans T., Steyaert J., Schiller PW., Manglik A., von Zastrow M. A Genetically Encoded Biosensor Reveals Location Bias of Opioid Drug Action. Neuron 2018;98(5):963–976.e5. Doi: 10.1016/j.neuron.2018.04.021.

139. Sengmany K., Hellyer SD., Albold S., Wang T., Jeffrey Conn P., May LT., Christopoulos A., Leach K., Gregory KJ. Kinetic and system bias as drivers of metabotropic glutamate receptor 5 allosteric modulator pharmacology. Neuropharmacology 2019;149:83–96. Doi: 10.1016/j.neuropharm.2019.02.005.

