## Supplementary Information for "Targeting of subcellular metabotropic glutamate receptor 5 signaling to modulate pain transmission"

^1^Drug Discovery Biology Theme, Monash Institute of Pharmaceutical Sciences, Monash University, Melbourne, Australia. ^2^Department of Chemistry and Biology, University of Santiago Chile, Santiago, CL. ^3^Department of Molecular Pathobiology, New York University College of Dentistry, New York, USA. ^4^School of Clinical Medicine, Faculty of Medicine, University of Queensland, Brisbane, Australia. ^5^Drug Delivery, Disposition and Dynamics Theme, Monash Institute of Pharmaceutical Sciences, Monash University, Melbourne, Australia. ^6^ARC Centre for Cryo-electron Microscopy of Membrane Proteins, Monash Institute of Pharmaceutical Sciences, Monash University, Parkville, VIC 3052, Australia. ^7^Department of Physiology, Monash Biomedicine Discovery Institute, Monash University, Melbourne, Australia.

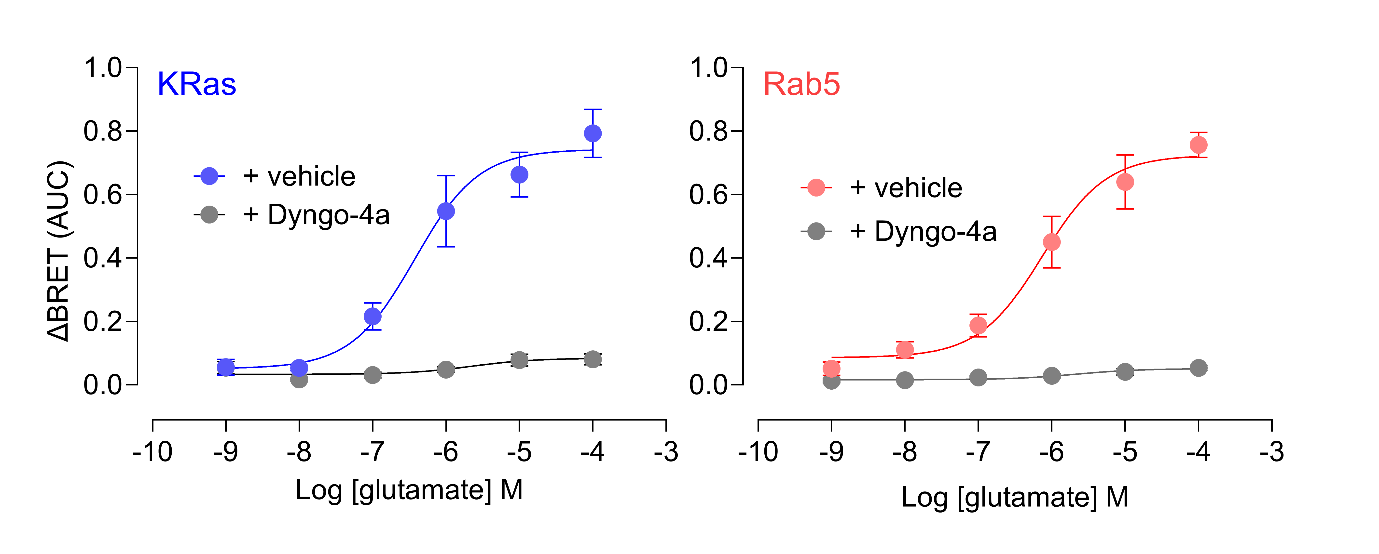

**Supplementary Figure 1**. **mGlu_5_ internalizes to early endosomes in a dynamin dependent manner.** Concentration response curves from BRET experiments for glutamate-induced movement of mGlu_5_ away from the plasma membrane (KRas, blue) and towards early endosomes (Rab5, red), expressed as AUC of kinetic BRET traces. Grey symbols indicate preincubation with the dynamin inhibitor Dyngo4a (30µM) for each marker. EC_50_ and E_max_ estimates are shown in Figure 1. Data are presented as mean ± s.e.m., n = 4-6 independent experiments performed in duplicate.

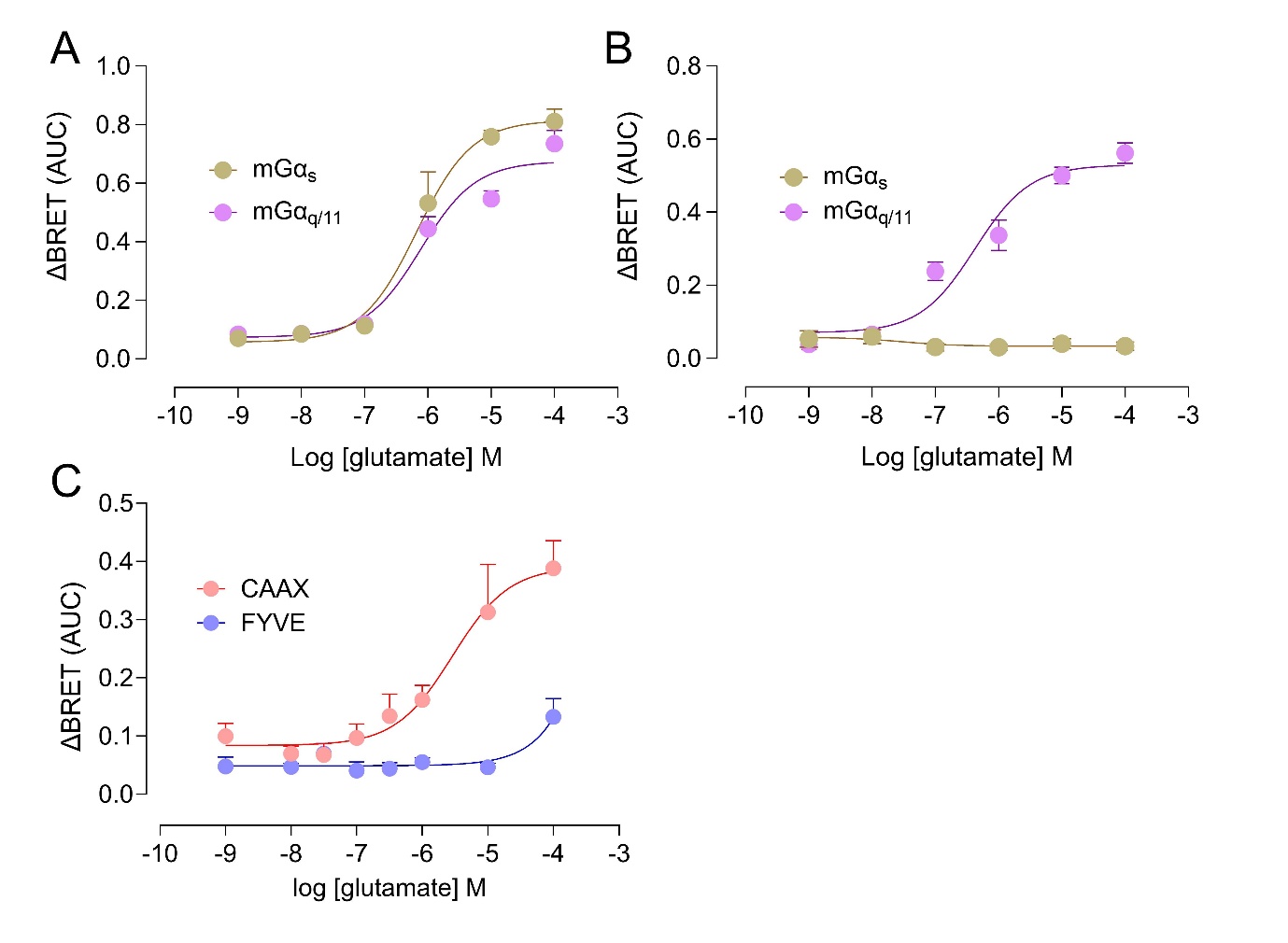

**Supplementary Figure 2.** **mGlu_5_ recruits mini-Gα_q_** **to both the cell surface and endosomes upon activation A)** Concentration-response curves for glutamate-induced G protein recruitment to mGlu_5_, expressed as AUC of kinetic BRET traces for mGlu_5-RLuc8_ association with mGα_s-Venus_ (brown) or mGαq_/11-Venus_ (purple) upon glutamate addition. **B**) Concentration-response curves for glutamate-induced G protein recruitment to endosomes, expressed as AUC of kinetic BRET traces for Rab5_-Venus_ association with mGα_s-RLuc8_ (brown) or mGα_q/11-RLuc8_ (purple) upon glutamate addition. **C)** Concentration-response curves for glutamate induced G protein recruitment to mGlu_5_ on the cell surface or in early endosomes, expressed as AUC of kinetic BRET traces for mGα_q/11-Venus_ association with mGlu_5-NP_ + CAAX-LgBit (cell surface, red) or mGlu_5-NP_ + FYVE-LgBit (early endosome, blue) upon glutamate addition. Data are presented as mean ± s.e.m., n = 4-6 independent experiments performed in duplicate.

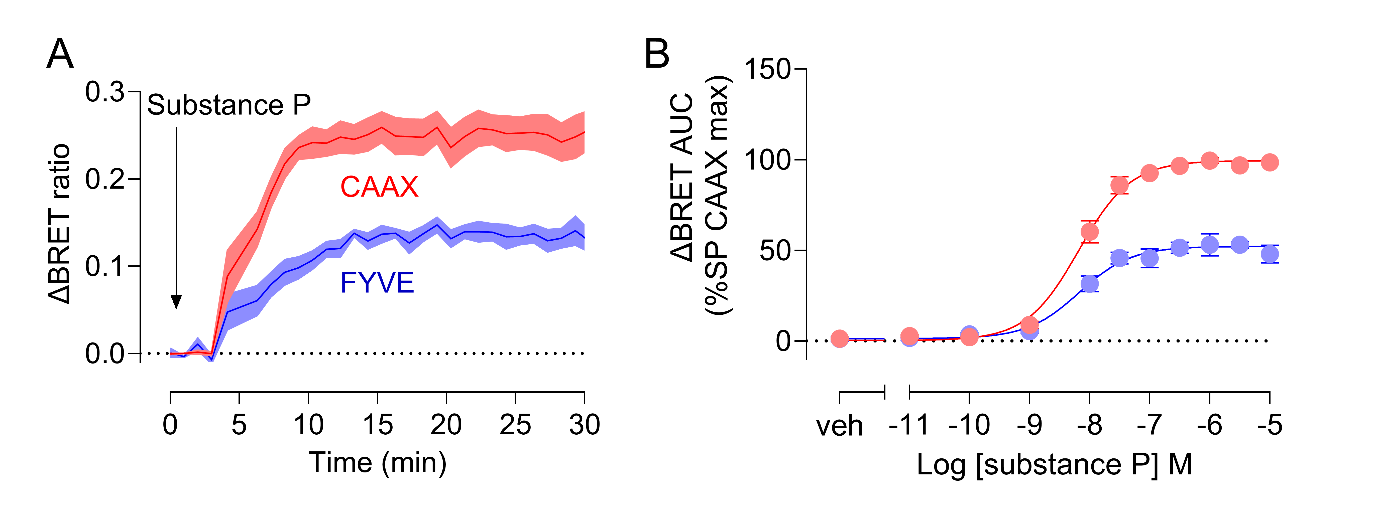

**Supplementary Figure 3**. **Validation of nanoBit constructs used in the current study.** HEK293A cells were co-transfected with mGα_q-Venus_, NK_1_R_-NP_ and CAAX-LgBit or FYVE-LgBit to determine G protein coupling at the plasma membrane and early endosomes, respectively. **A**) Kinetic traces of G protein coupling at the plasma membrane (CAAX, red) or early endosomes (FYVE, blue) after addition of substance P (30nM). **B)** Concentration-response curves for substance P-induced G protein coupling to NK_1_R at the plasma membrane (red) and early endosomes (blue). Data are presented as mean ± s.e.m., n = 5 independent experiments performed in duplicate.

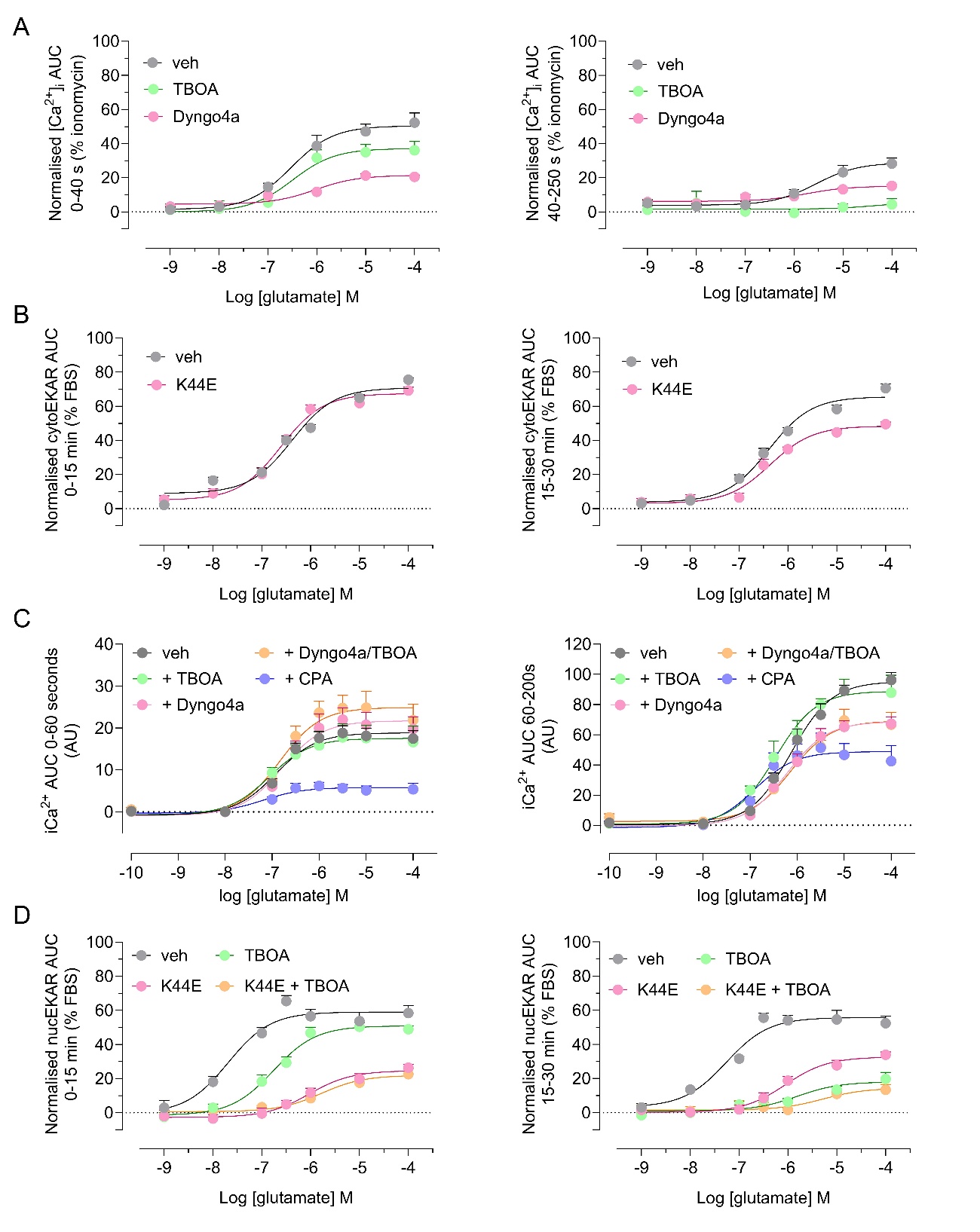

**Supplementary Figure 4. Sustained mGlu_5_ signalling relies on receptor internalization and glutamate transport.** Concentration-response curves for glutamate signalling in global iCa^2+^ mobilization (**A**), cytosolic ERK1/2 phosphorylation (cytoEKAR) (**B**), nuclear iCa^2+^ mobilization (**C**) and nuclear ERK1/1 phosphorylation (nucEKAR) (**D**) assays under different conditions: vehicle (black), dynamin inhibition (Dyngo-4A, 30µM; or K44E co-transfection, pink), glutamate transport inhibition (DL-TBOA, 50µM, green), combined dynamin-glutamate transport inhibition (orange) or SERCA pump inhibition (CPA, 30µM, blue). Kinetic traces were divided into first and second phase for each assay, presented as left and right panels, respectively. Data are presented as mean + s.e.m., n = 4-6 independent experiments performed in duplicate.

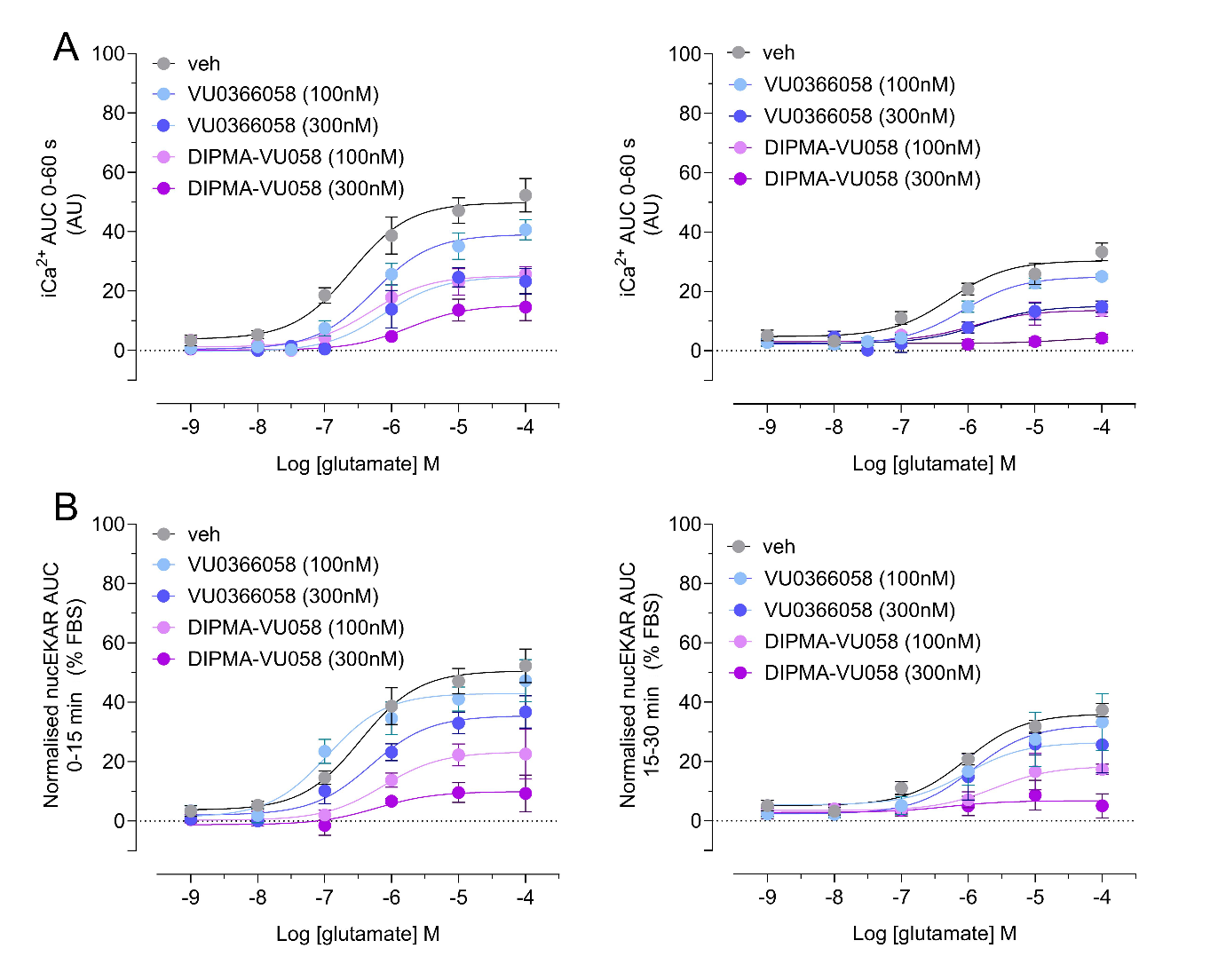
**Supplementary Figure 5. Nanoparticle encapsulation enhances VU0366058 inhibition of glutamate-induced cytosolic iCa^2+^and nuclear ERK1/2 signalling in HEK293A cells**. Concentration-response curves for glutamate-induced global iCa^2+^ mobilization (**A**) or nuclear ERK1/2 signalling (**B**) in the absence (black) and presence of VU0366058 (blue) or DIPMA-VU058 (purple). Cells were preincubated with the indicated concentrations of VU0366058 or DIPMA-VU058 for 30 min prior to glutamate addition. Data are expressed as AUC of kinetic traces. Data are presented as mean ± s.e.m., n = 4-6 independent experiments performed in duplicate.
